# MK2/p38/p53 suppress basal IL-1β and non-canonical NF-κB signaling

**DOI:** 10.1101/2025.09.30.679163

**Authors:** Sarah M. Herr, Diana Stalkopf, Sofie Padaszus, Lukas A. Herbst, Anneke Dörrie, Rainer Niedenthal, Natalia Ronkina, Tatiana Yakovleva, Alexey Kotlyarov, Matthias Gaestel

## Abstract

Interleukin (IL)-1β is a pro-inflammatory cytokine implicated in sterile inflammation and tumor development. Investigating the role of MAPKAP kinase 2 (MK2) in IL-1β processing, we found that *Il1b* mRNA and IL-1β protein levels were elevated in resting *MK2*-knockout (KO) macrophages and in the serum of *MK2/3*-double-KO mice. This was linked to activation of the non-canonical NF-κB pathway in the absence of MK2 or its activator, p38α. Rescue by MK2, its kinase-inactive mutant MK2K79R, or p38α suppressed this pathway and reduced *Il1b* expression. We also observed decreased basal protein levels of tumor suppressor p53 in *MK2*-or *p38α*-deficient cells. Mechanistically, p53 interacts with mitochondrial caspase-3, promoting cleavage of RelB, thereby inhibiting non-canonical NF-κB signaling and subsequent *Il1b* and *TP53* expression. These findings explain elevated basal IL-1β levels in *MK2*-KO macrophages and uncover a new autoregulatory mechanism of *TP53* expression. Additionally, they reveal a new mechanism that contributes to the long-discussed link between cancer and inflammation, wherein the tumor suppressor p53 inhibits cytokine production in parallel.

## Introduction

p38 mitogen-activated protein kinases (p38^MAPK^) comprise four distinct isoforms - p38α (MAPK14), p38β (MAPK11), p38γ (MAPK12 or ERK6), and p38δ (MAPK13 or SAPK4) - that are part of highly conserved signaling cascades. They are activated by various cellular stresses, such as inflammatory cytokines, growth factors, bacterial lipopolysaccharides (LPS), osmotic stress, and ultraviolet irradiation. p38 signaling affects gene expression at the levels of transcription, mRNA stability, and translation. It also regulates survival, apoptosis, differentiation, aggregation, and migration (*1*). MK2 and MK3 are the only MAPK-activated protein kinases (MKs) that are exclusively phosphorylated and activated by p38α/β. MK2 and MK3 are closely related in terms of structure and function and share the same substrates. However, MK2 is expressed and active at higher levels than MK3 in most cells and tissues (*2*). p38α/β binds to a docking motif in the C-terminus of MK2 and phosphorylates MK2 regulatory sites in response to stress. Consequently, MK2 and p38 are co-transported from the nucleus to the cytoplasm, where MK2 substrates are phosphorylated (*3*). MK2 has several specific substrates, including tristetraprolin (TTP), serum response factor (SRF), heat shock proteins Hsp25/27, receptor-interacting serine/threonine-protein kinase 1 (RIPK1), and RNA-binding motif protein 7 (RBM7). These substrates are involved in the regulation of immediate early gene responses, cytokine expression, cell death and migration (*4–8*). Additionally, MK2 stabilizes p38 independently of its catalytic activity through the formation of a binary protein complex (*3*).

p53 is a transcription factor that is responsible for regulating several genes involved in the cell cycle control, apoptosis, and senescence. It also exhibits transcription-factor-independent functions, particularly in the regulation of apoptosis through the mitochondrial pathway. Its medical significance is highlighted by the fact that p53 mutations are present in over 50% of human tumors (*9*). In resting cells, p53 is constitutively expressed, but becomes ubiquitinated by Mouse double minute 2 homolog (MDM2) and subsequently degraded by the proteasomes, maintaining low protein levels. Following exposure to DNA-damaging agents or other stress stimuli, p53 undergoes a series of post-translational modifications that stabilize and activate p53 by reducing its interaction with MDM2. These modifications include the phosphorylation of Ser^15^ and Ser^37^ (*10, 11*). The stabilization of p53 is also adjusted by a broad variety of other interacting proteins. p38 exists in a physical complex with the tumor suppressor p53 and coexpression of p38 stabilizes p53 protein. After UV radiation, p38 phosphorylates p53 at Ser^33^ and Ser^46^. These sites are associated with p53-mediated apoptosis and are important for the subsequent phosphorylation of Ser^15^ and Ser^37^ of p53 (*12*).

The transcription factor NF-κB plays a crucial role in regulating the immune responses, cell proliferation and cell death. There are five known NF-κB proteins: NF-κB1 (p105/p50), NF-κB2 (p100/p52), RelA (p65), RelB, and c-Rel. These proteins are classified into the canonical and the non-canonical NF-κB pathways. The canonical NF-κB pathway is triggered by a wide variety of receptors that activate TGF-β-activated kinase 1 (TAK1). Then, TAK1 mediates the phosphorylation of the inhibitor of NF-κB kinase (IKK) IKKβ. The activated IKK complex, composed of IKKα, IKKβ and IKKγ/Nuclear Factor-κB-Essential Modulator (NEMO) phosphorylates the inhibitor of κB (IκB) IκBα and the IκB-like molecule p105, which subsequently are ubiquitinated and degraded. The cleavage product of the NF-κB1 precursor protein p105, p50, is translocated to the nucleus as a homodimer or a heterodimer together with RelA or c-Rel. There, they bind to specific DNA elements to induce the transcription of canonical target genes. In contrast, the non-canonical NF-κB pathway is activated only by a subset of TNF receptor superfamily members. Under resting conditions, NF-κB-inducing kinase (NIK) is recruited to an ubiquitin ligase complex consisting of a cellular inhibitor of apoptosis (cIAP), tumor necrosis factor (TNF)-receptor-associated factors (TRAF)-2 and TRAF-3. This leads to the constitutive degradation of NIK. After stimulation, TRAF3 is degraded, resulting in the release and stabilization of NIK. NIK then phosphorylates IKKα, which in turn phosphorylates the NF-κB2 precursor protein p100. Subsequently, the cleavage product p52 and RelB are then translocated into the nucleus, where they activate target genes of the non-canonical NF-κB pathway (*13*).

Here we demonstrate, that in the absence of MK2 or p38α, basal p53 protein level is reduced and in turn, the non-canonical NF-κB pathway is activated. Subsequently, the mRNAs of pro-inflammatory interleukin IL-1β coding gene *Il1b* as well as the p53 coding gene *TP53* are enriched. Accumulation of p53 activates mitochondrial caspase-3, which cleaves RelB. RelB is a direct transcription factor of p53 (*14*). Subsequently, the non-canonical NF-κB pathway and *TP53* transcription itself are inactivated, indicating a novel autoregulatory mechanism of p53 gene expression.

## Results

### *MK2/3*-DKO mice display elevated levels of IL-1β

To investigate whether the kinases MK2 and MK3 regulate IL-1β, the *Il1b* mRNA level was analyzed in bone marrow-derived macrophages (BMDMs) from wild-type (WT) and *MK2/3*-DKO mice that were either left untreated or stimulated with either IL-1α or LPS. IL-1α is usually released by dying cells, necrotic cells or stressed cells and occurs in sterile chronic diseases, while LPS is derived from bacteria. The expression and release of IL-1β typically occurs in response to two distinct signals. IL-1α or LPS serves as the first signal, resulting in the expression of *Il1b*. The *Il1b* mRNA levels were already elevated in unstimulated BMDMs of the *MK2/3*-DKO (Fig. 1A), which increased further after IL-1α stimulation (Fig. 1B). The *Il1b* mRNA levels in LPS-treated BMDMs were strongly increased, but did not differ between the different genotypes (Suppl. 1A). Consistent with the elevated mRNA levels, *MK2/3*-DKO cells also displayed elevated IL-1β protein levels in unstimulated BMDMs (Fig. 1C). Usually, a second signal, such as ATP or nigericin, is required for activation of the inflammasome complex, caspase-1, and the subsequent release of IL-1β. Even without a second stimulus, the concentration of IL-1β in the supernatant of unstimulated *MK2/3*-DKO-BMDMs was significantly higher than that in WT-BMDMs (Fig. 1D), whereas the cell viability was not affected (Suppl. 1B). After the addition of the second stimulus, such as nigericin or ATP, *MK2/3*-DKO BMDMs pretreated with IL-1α released significantly higher levels of IL-1β compared to WT BMDMs (Fig. 1 E, F). In contrast, equal concentrations of IL-1β were released after the addition of ATP to LPS-pretreated cells (Suppl. 1C). Analysis of IL-1 receptor mRNA expression (*IL-1r1*, *IL-1r2*, and *IL-1rn*) revealed no differences among WT and *MK2/3*-DKO BMDMs (Suppl. 1D-F).

**Figure 1:**
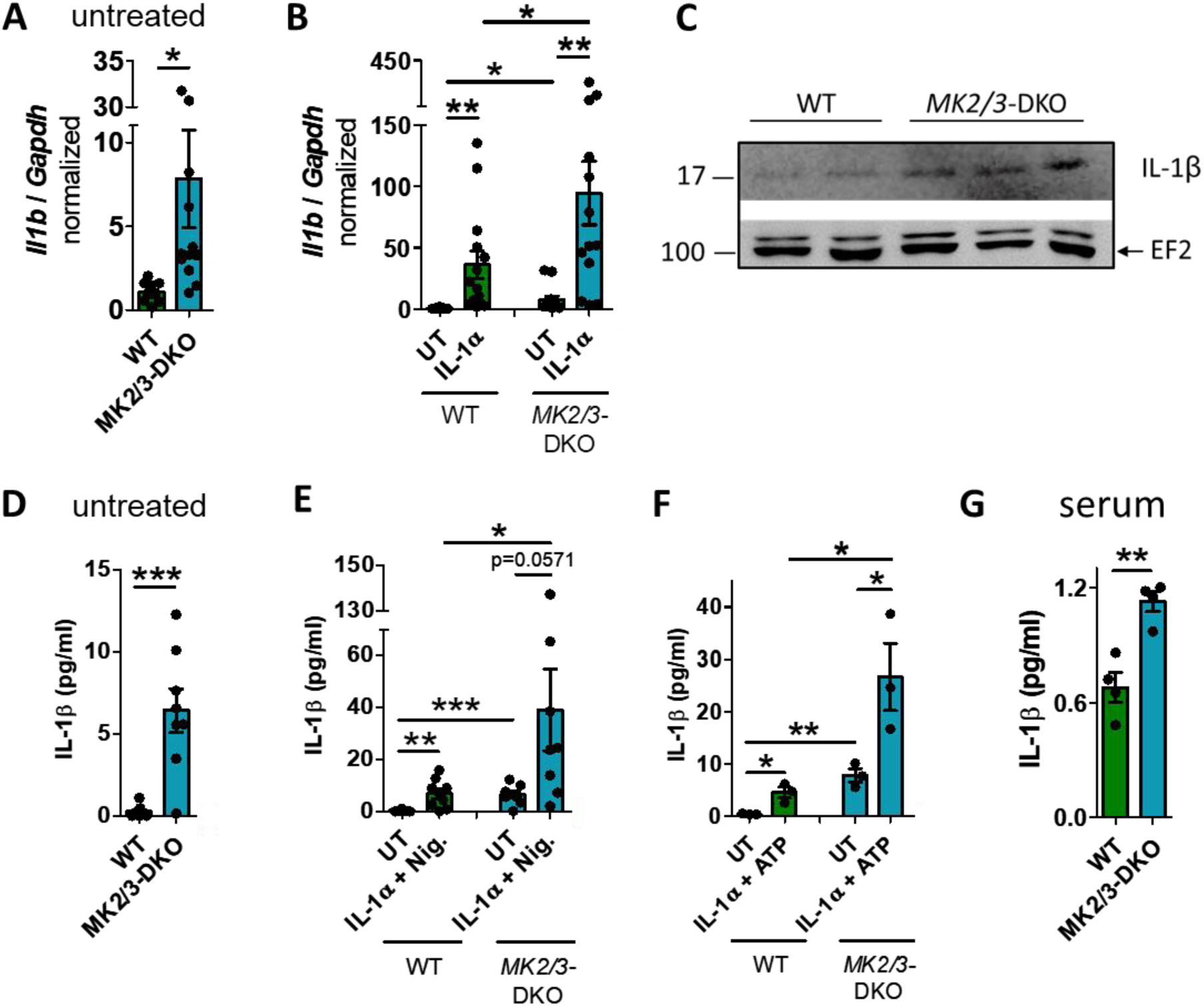
Interleukin (IL)-1β levels are elevated in *MK2/3* double knockout (DKO) mice. *Il1b* mRNA levels are increased in **(A)** untreated and **(B)** IL-1α-treated (5 ng/ml, 1h) *MK2/3*-DKO bone marrow-derived macrophages (BMDMs) compared to wild type (WT). WT n=14, DKO n=13. **(C)** Basal IL-1β protein levels are elevated in *MK2/3*-DKO BMDM. Elongation factor 2 (EF2) serves as a control. One representative Western Blot of total WT n=4, DKO n=6. **(D)** The concentration of IL-1β is higher in the supernatant of untreated and **(E)** IL-1α (5ng/ml, 4h) +Nigericin (20 µM, 8h) or **(F)** IL-1α+ATP (5 mM, 5.5h)-treated *MK2/3*-DKO BMDM than in WT. D-E) WT n=9, DKO n=8. F) n=3/group. **(G)** The basal concentration of IL-1β is elevated in the serum of *MK2/3*-double knockout (DKO) mice. n=6 mice/group, whereby one sample/group was pooled from 3 mouse sera. Mean ± SEM, students t-test, * P<0.05, ** P<0.01, *** P<0.001.

Since unstimulated BMDMs from *MK2/3*-DKO mice exhibit elevated *Il1b* mRNA and IL-1β protein levels and release more IL-1β than unstimulated BMDMs from WT mice, we analyzed the sera of *MK2/3*-DKO and WT mice from the same breeding to determine their basal cytokine levels. *MK2/3*-DKO serum showed significantly higher basal IL-1β levels than WT serum (Fig. 1G), demonstrating the relevance in vivo. However, there was no significant difference in the basal levels of TNF-α and CXCL-1 (Suppl. 1G-H).

### MK2 and MK3 suppress the basal and IL-1α-induced IL-1β level in rescued immortalized *MK2*-KO BMDMs

Next, *MK2*-KO BMDMs were immortalized and transduced with either *MK2* (i*MK2*-KO+*MK2*) or with an empty vector (i*MK2*-KO+*GFP*) as control (Suppl. 1I). Rescuing *MK2* suppressed the elevated *Il1b* mRNA level (Fig. 2A) to a level comparable to the WT BMDMs (Fig. 1A, 2A). In contrast, the basal *Tnf* mRNA level was unaffected in i*MK2*-KO cells, resembling the situation in WT BMDMs (Suppl. 1J). IL-1α treatment increased the differences in the *Il1b* mRNA levels in i*MK2*-KO cells as well (Fig. 2B). This was confirmed in an additional cell line. RAW macrophages treated with *MK2* siRNA had higher *Il1b* mRNA levels compared to the control after IL-1α stimulation (Fig. 2C, Suppl. 2A). To investigate, whether MK3 is functionally redundant with MK2, i*MK2*-KO cells were rescued with *MK3*. Indeed, MK3 also repressed the *Il1b* mRNA level in both unstimulated and IL-1α-treated cells (Fig. 2D, E), as well as the basal IL-1β protein level (Fig. 2F). Therefore, MK2 and MK3 redundantly suppress IL-1β production in these cells.

**Figure 2:**
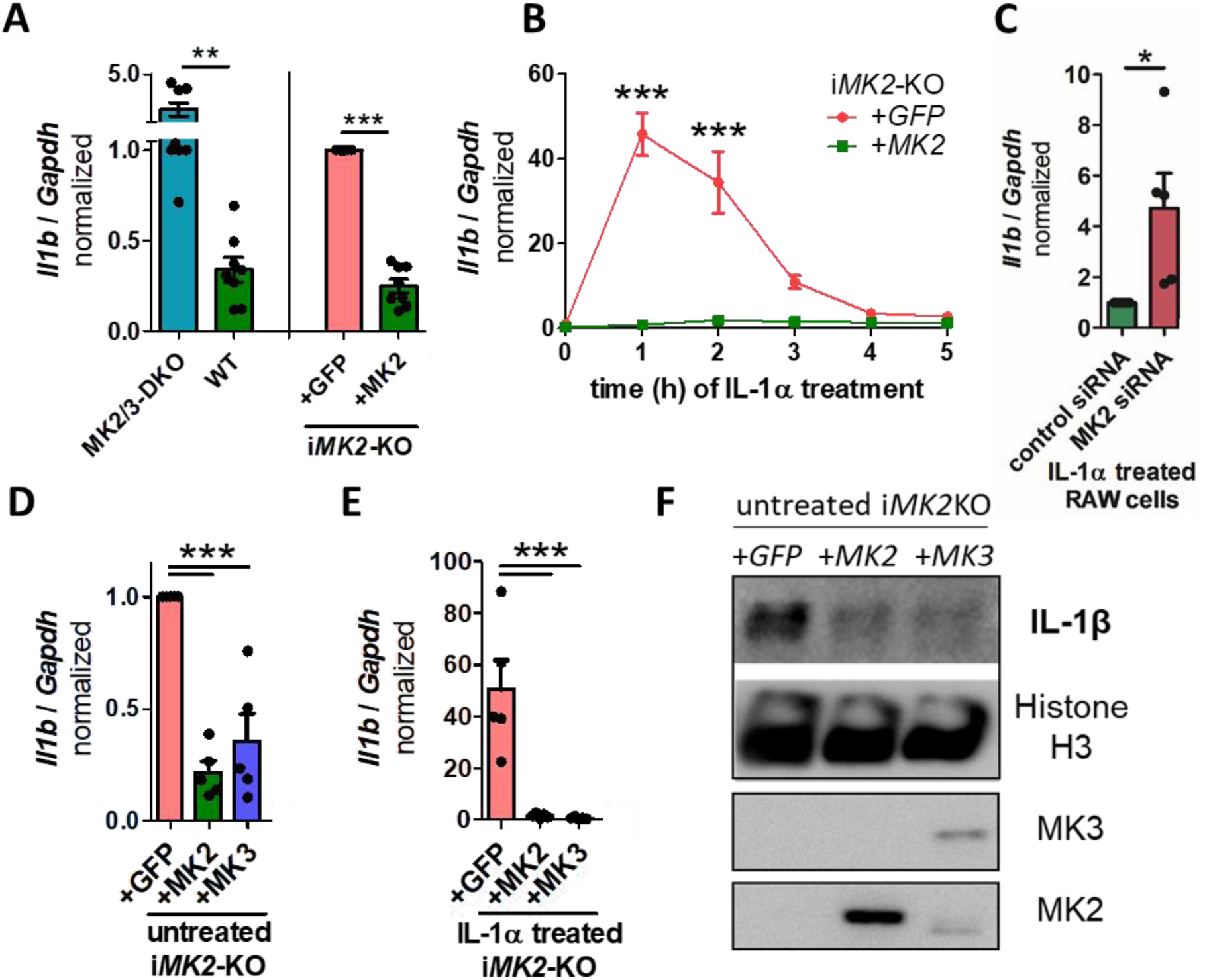
The level of IL-1β is increased in immortalized *MK2*-KO (i*MK2*-KO) cells. **(A)** i*MK2*-KO cells transduced with an empty vector (+*GFP*) showed elevated levels of *Il1b* mRNA compared to *MK2*-rescued cells (+*MK2*) (right, n=8), similar to those observed in *MK2/3*-DKO and WT bone-marrow-derived macrophages (BMDMs) (left; WT n=9, DKO n=8). **(B)** i*MK2*-KO+*GFP* cells show increased *Il1b* mRNA after IL-1α treatment (5 ng/ml) compared to *MK2*-rescued cells. n=3. **(C)** RAW cells treated with *MK2* siRNA have higher *Il1b* mRNA levels compared to the control after IL-1α stimulation. **(D)** Similar to *MK2*, rescuing *MK3* decreases the level of *Il1b* mRNA in resting or **(E)** IL-1α (5 ng/ml, 1h)-treated i*MK2*-KO cells, **(F)** as well as the basal level of IL-1β protein. Histone H3 serves as a control. **(A, C)** students t-test, **B)** 2W-RM-ANOVA with Bonferroni posttests, **(D, E)** 1W-ANOVA with Tukey’s Multiple Comparison Test, mean ± SEM, ** P<0.01, *** P<0.001.

### MK2 suppresses the non-canonical NF-κB pathway

To identify the pathway activated in the absence of MK2, several inhibitors were tested. Actinomycin D, an inhibitor of RNA transcription, was used as a positive control and efficiently suppressed the basal *Il1b* mRNA level (Suppl. 2B). cFos/AP1 and TRAF6 inhibitors did not affect the *Il1b* mRNA levels in i*MK2*-KO cells (Suppl. 2C, D). Suppressing the canonical NF-κB pathway with three specific IKKβ inhibitors (Takinib, sc-514, CAY10657) did not reduce the basal *Il1b* mRNA level in i*MK2*-KO cells. However, they suppressed IL-1α- and LPS-induced *Il1b* mRNA levels (Fig. 3A, Suppl. 2E). In contrast, the simultaneous inhibition of IKKα and IKKβ with HPN-01 also inhibited the basal *Il1b* mRNA expression (Fig. 3A). These results were confirmed with BMS 345541. At low concentrations, BMS 345541 solely inhibits IKKβ, resulting in no reduction of the basal *Il1b* mRNA level. At higher concentrations, where both IKKα and IKKβ are suppressed simultaneously, the basal *Il1b* mRNA level was significantly reduced in i*MK2*-KO cells (Suppl. 2F). Consequently, the specific inhibition of the non-canonical NIK/ IKKα by B022 decreased in particular the basal *Il1b* mRNA and *Map3k14* mRNA expression in i*MK2*-KO cells (Fig. 3A, Suppl. 2G). Inhibition of the non-canonical NF-κB pathway by B022 reduced also basal IL-1β protein level in i*MK2-*KO cells (Fig. 3B, C) as well as basal *Il1b* mRNA in primary *MK2/3*-DKO BMDMs (Fig. 3D).

**Figure 3:**
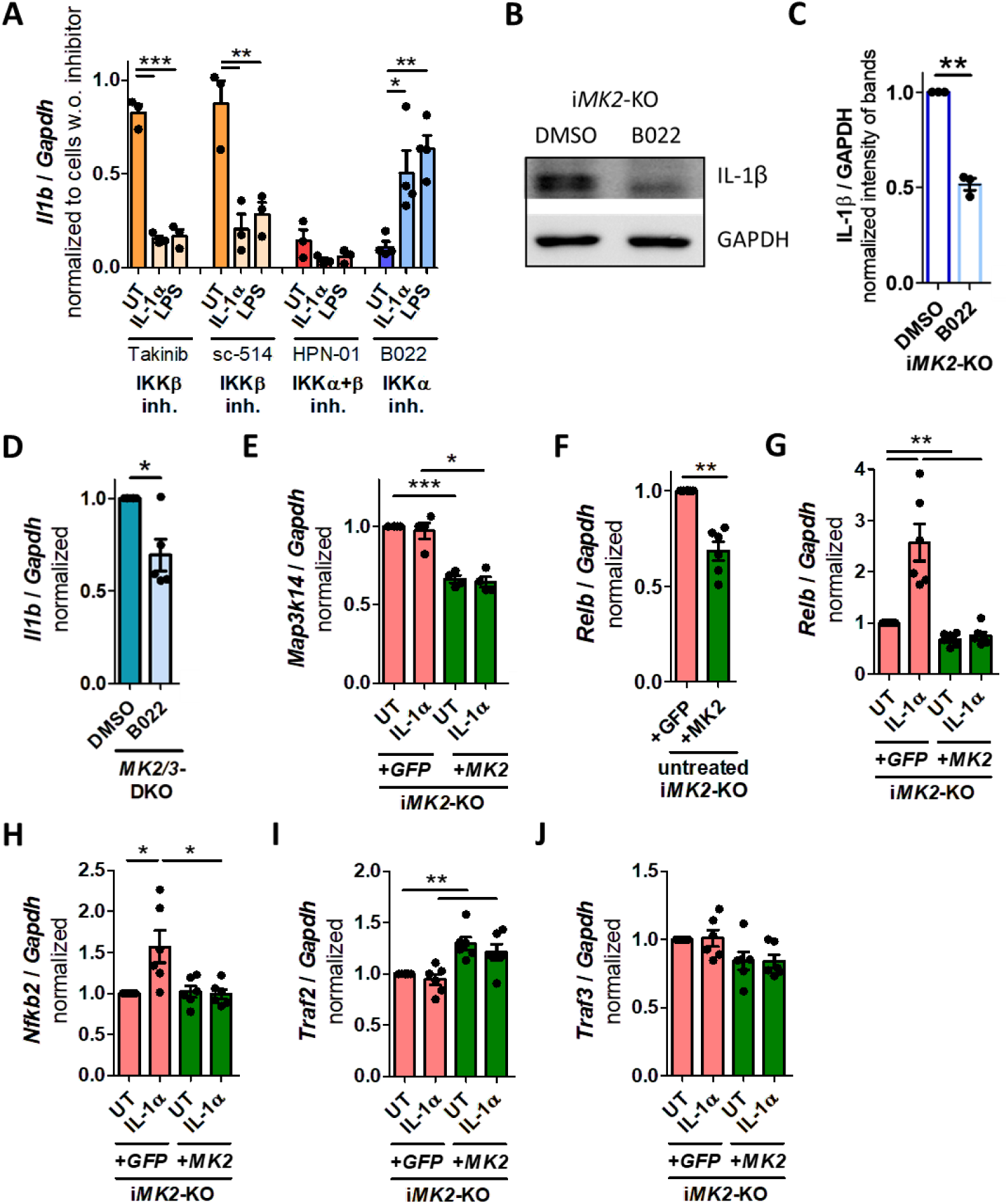
The non-canonical NF-κB pathway is activated in i*MK2*-KO cells (mRNA) **(A)** Inhibition of the canonical NF-κB pathway using the IKKβ inhibitors Takinib (10 µM, 2h) and sc-514 (50 µM, 2h) reduced *Il1b* mRNA levels in IL-1α- and LPS-treated i*MK2*-KO cells, but did not affect basal *Il1b* levels. Inhibiting IKKα and IKKβ with HPN-01 (10 µM, 2h) reduced *Il1b* mRNA levels in untreated (UT) and IL-1α (5 ng/ml, 1h)- or LPS-stimulated cells (100 ng/ml, 1h). Inhibition of the non-canonical NF-κB pathway by IKKα inhibitor B022 (5 µM, 2h) reduced mainly basal *Il1b* mRNA. **(B, C)** IL-1β protein level is reduced in resting i*MK2*-KO cells after treatment with B022 (5 µM, 7h). GAPDH serves as a control. **(D)** *Il1b* mRNA is reduced in *MK2/3*-DKO BMDMs after treatment with B022 (5 µM, 2h). **(E)** The level of *Map3k14* mRNA is increased in i*MK2*-KO+*GFP*. **(F)** *Relb* mRNA level is increased in UT i*MK2*-KO+*GFP* cells. **(G)** *Relb* and **(H)** *Nfkb2* mRNA levels are increased in IL-1α (5ng/ml, 1h)-stimulated i*MK2*-KO+*GFP* cells. **(I)** Basal *Traf2* mRNA is reduced in i*MK2*-KO+*GFP* cells. **(J)** The *Traf3* mRNA level is not changed significantly. **(A)** 1W-ANOVA with Tukey’s Multiple Comparison Test, **(C-J)** students t-test, mean ± SEM, * P<0.05, ** P<0.01, *** P<0.001.

i*MK2*-KO cells displayed higher basal mRNA expression levels of *Map3k14*, the NIK-coding gene that activates the non-canonical NF-κB pathway (Fig. 3E). Additionally, i*MK2*-KO cells also showed elevated basal levels of the mRNA of non-canonical *Relb* (Fig. 3F). *Relb* and *Nfkb2* mRNA expression increased after IL-1α treatment solely in i*MK2*-KO+*GFP* cells, but not in MK2-rescued i*MK2*-KO+*MK2* cells (Fig. 3G, H). There was no significant difference in canonical *Nfkb1* and *Rela* mRNA expression (Suppl. 2H, I). Furthermore, *Traf2* and *Traf3* were analyzed, as both are known negative regulators of NIK (*13, 15*). The basal level of *Traf2*, but not *Traf3*, mRNA was significantly reduced in i*MK2*-KO macrophages (Fig. 3I, J).

Next, we analyzed the levels of signaling proteins in the nuclear and cytoplasmic fractions of i*MK2*-KO+*GFP* and i*MK2*-KO+*MK2* BMDMs using Western blotting. Following IL-1α treatment, the nuclear fraction of the i*MK2*-KO+*GFP* cells exhibited higher levels of the non-canonical proteins RelB and NF-κB2, as well as the canonical proteins RelA and NF-κB1 (Fig. 4A). Notably, only the non-canonical NF-κB pathway proteins RelB and NF-κB2 were elevated in the nuclear and cytoplasmic fractions of unstimulated i*MK2*-KO+*GFP* cells, as compared to the same fractions of i*MK2*-KO+*MK2* cells. No differences in basal protein levels were detected among the members of the canonical NF-κB pathway: RelA, NF-κB1, and IκBα (Fig. 4A-C, Suppl.3A-C). Furthermore, whole-cell lysis revealed decreased TRAF2 protein levels in unstimulated i*MK2*-KO cells. However, the basal level of TRAF3 and cIAP remained unaffected in the absence of MK2. Whole-cell lysis also confirmed an overall reduction of RelB in the presence of MK2 (Fig. 4D, E, Suppl.3D). Since TRAF2 mediates the proteasome-dependent degradation of the pro-inflammatory transcription factor c-Rel (*16*), we also analyzed c-Rel protein expression. Indeed, i*MK2*-KO+*GFP* cells had higher c-Rel protein levels in the nucleus and cytoplasm than in i*MK2*-KO+*MK2* cells (Fig. 4F, Suppl. 3E).

**Figure 4:**
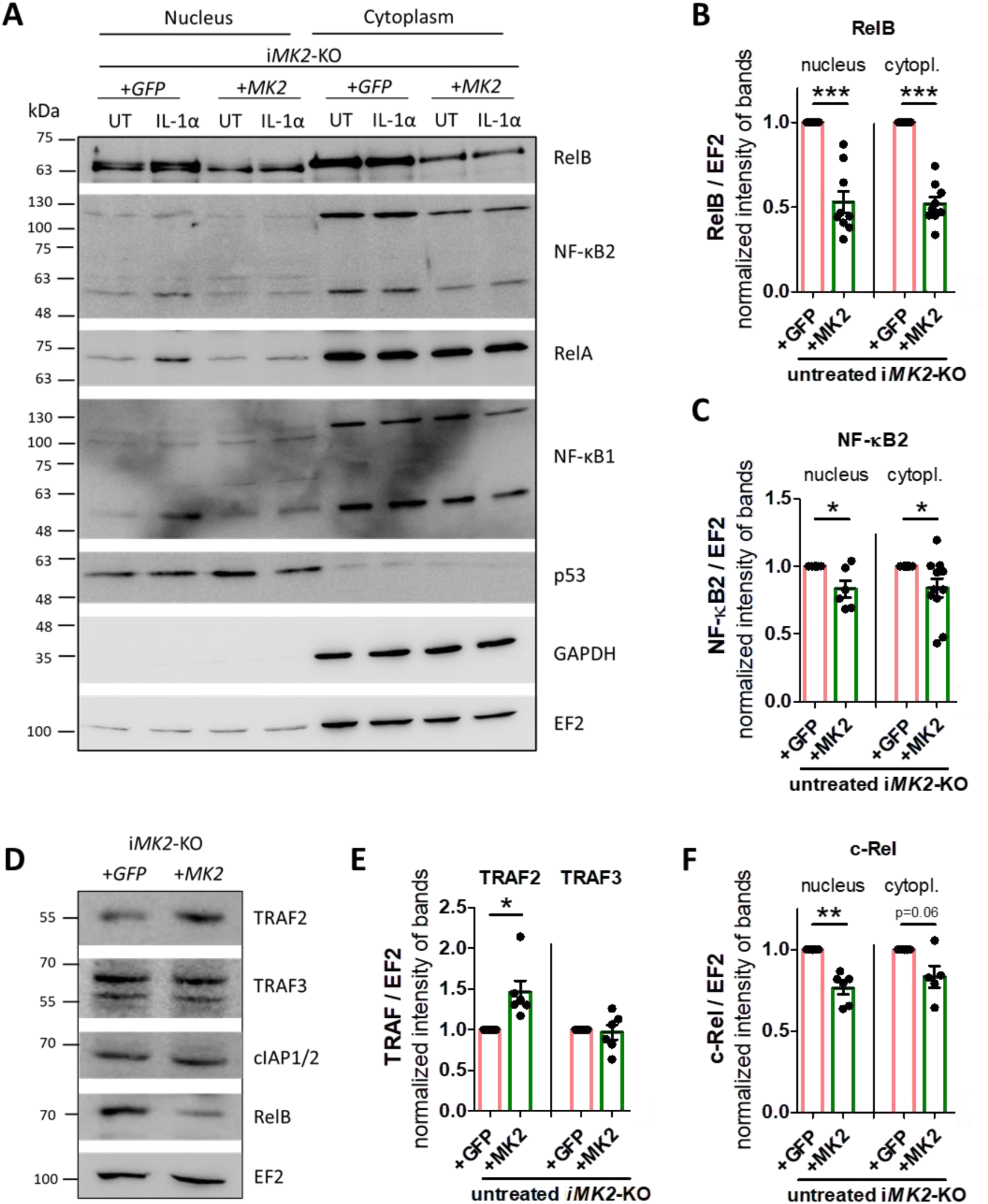
The non-canonical NF-κB pathway is activated in i*MK2*-KO cells (protein) **(A-C)** Compared to i*MK2*-KO+*MK2* cells, untreated (UT) i*MK2*-KO+*GFP* cells have higher levels of the non-canonical proteins RelB and NF-κB2 in nuclear and cytoplasmic fractions, but not of the canonical RelA and NF-κB1 proteins. Following IL-1α treatment (5 ng/ml, 2h), the nuclear fraction of i*MK2*-KO cells showed elevated protein levels of RelB, NF-κB2, RelA, and NF-κB1. p53 and GAPDH serve as controls for successful nuclear/cytoplasmic separation. EF2 acts as a general control, used for normalization. **(D-E)** The basal protein level of TRAF2 is reduced in whole cell lysis of i*MK2*-KO+*GFP* cells, whereas TRAF3 and cIAP1/2 are not changed. **(F)** The protein level of c-Rel is increased in i*MK2*-KO+*GFP* cells. mean ± SEM, students t-test, * P<0.05, ** P<0.01, *** P<0.001.

RNA sequencing of the immortalized BMDMs revealed increased mRNA levels of IL-1β pathway components (such as *Il1b*, *Il18*, and *Nlrp3*) and of non-canoncial NF-κB pathway components (such as *Chuk*, *Nfkb2*, and *Relb*), as well as a wide range of non-canonical NF-κB pathway targets predicted using known transcription factor target site motifs (such as *Birc2*, *Birc3*, *Icam1*, and *Rel*) (*17–19*) in i*MK2*-KO+*GFP* cells (Fig. 5).

**Figure 5:**
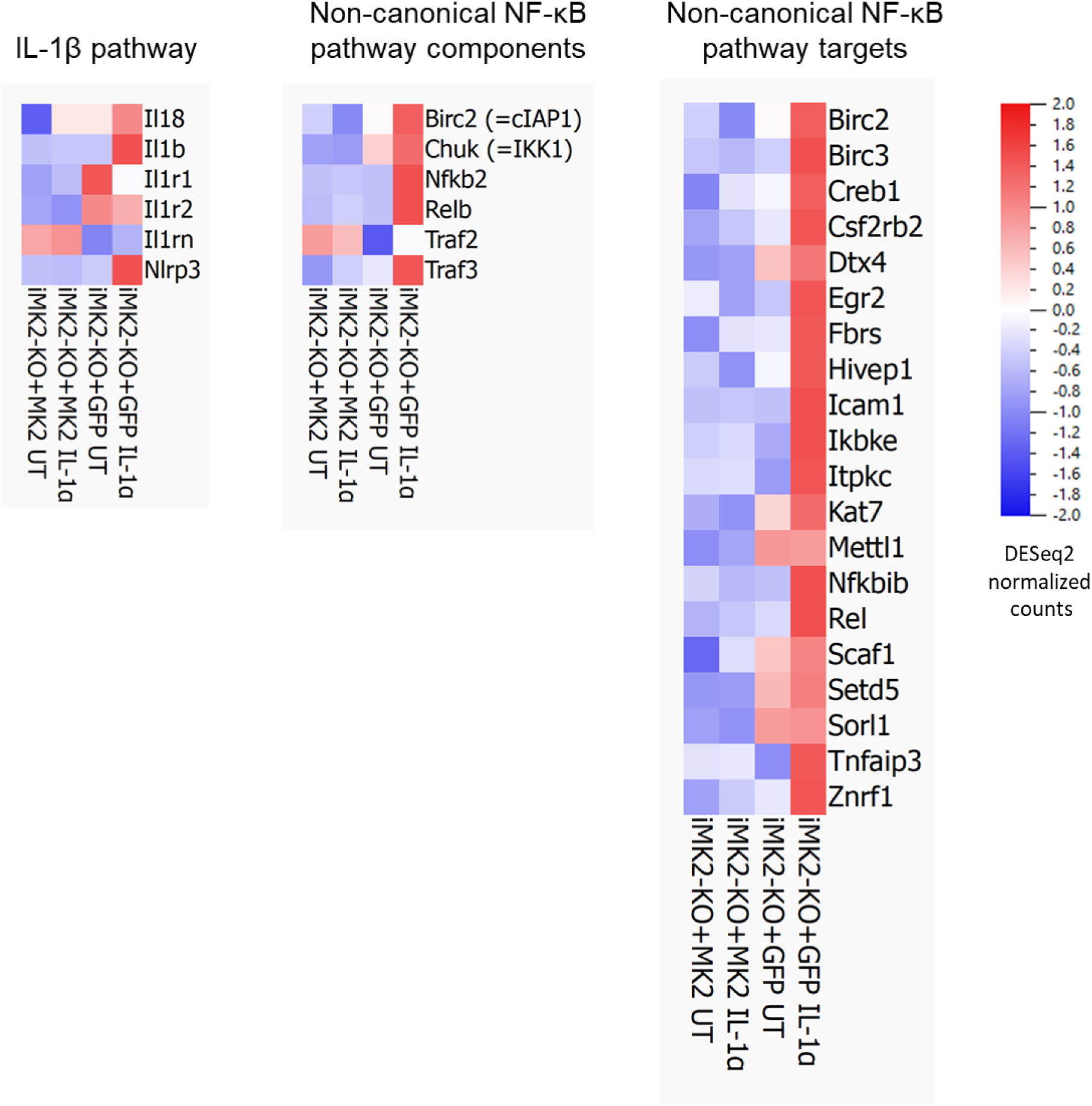
The non-canonical NF-κB pathway is activated in i*MK2*-KO cells. RNA sequencing revealed increased levels of mRNA for components of the IL-1β and non-canonical NF-κB pathways, as well as for targets of the non-canonical NF-κB pathway, in i*MK2*-KO+*GFP* cells. This finding was reinforced by IL-1α treatment (5 ng/ml, 1h).

*MK2*-KO macrophages exhibited elevated basal mRNA and protein levels of the non-canonical NF-κB pathway components NIK (*MAP3K14*), RelB, and NF-κB2. Conversely, they displayed decreased mRNA and protein levels of TRAF2, the negative regulator of NIK. Neither the basal mRNA nor basal protein levels of canonical components RelA, NF-κB1, or IκBα are changed in the absence of MK2. In addition, inhibition of the non-canonical, but not the canonical NF-κB pathway, reduced the basal *IL1b* mRNA level in i*MK2*-KO cells. Therefore, MK2 suppresses the upregulation of *IL1b* in unstimulated cells by blocking the non-canonical NF-κB pathway.

### Stabilization of the protein kinase p38α by MK2/3 is a mechanism for suppressing *Il1b*

To examine the role of MK2’s catalytic activity in suppressing *Il1b*, i*MK2*-KO cells were rescued using the kinase-inactive mutant *MK2K79R* (Suppl. 4A). Catalytic inactivation of MK2 did not affect the suppression of *Il1b* mRNA levels in unstimulated or IL-1α-treated cells (Fig. 6A-B). Beside of its catalytic activity, MK2 also binds and stabilizes its activator, the protein kinase p38α (*20–22*). Indeed, i*MK2*-KO+*GFP* BMDMs express significantly less endogenous p38α protein than i*MK2*-KO+*MK2* cells, whereas the *MAPK14* (p38α) mRNA level is not affected (Fig. 6C, Suppl. 4B-C). Because the C-terminal extension of MK2 is involved in binding to p38, a mutant lacking this extension (*MK2-Δ365-386*) was transduced into i*MK2*-KO cells that had a comparable level of MAPKAPK2 expression to MK2 (Suppl. 4D). Although this mutant is catalytically active, it could not restore the endogenous p38α protein level and was unable to suppress *Il1b* mRNA levels in untreated or IL-1α-treated cells (Fig. 6C-D, Suppl. 4E). These findings strongly suggest that MK2’s inability to stabilize p38 is the primary reason for *Il1b* upregulation.

**Figure 6:**
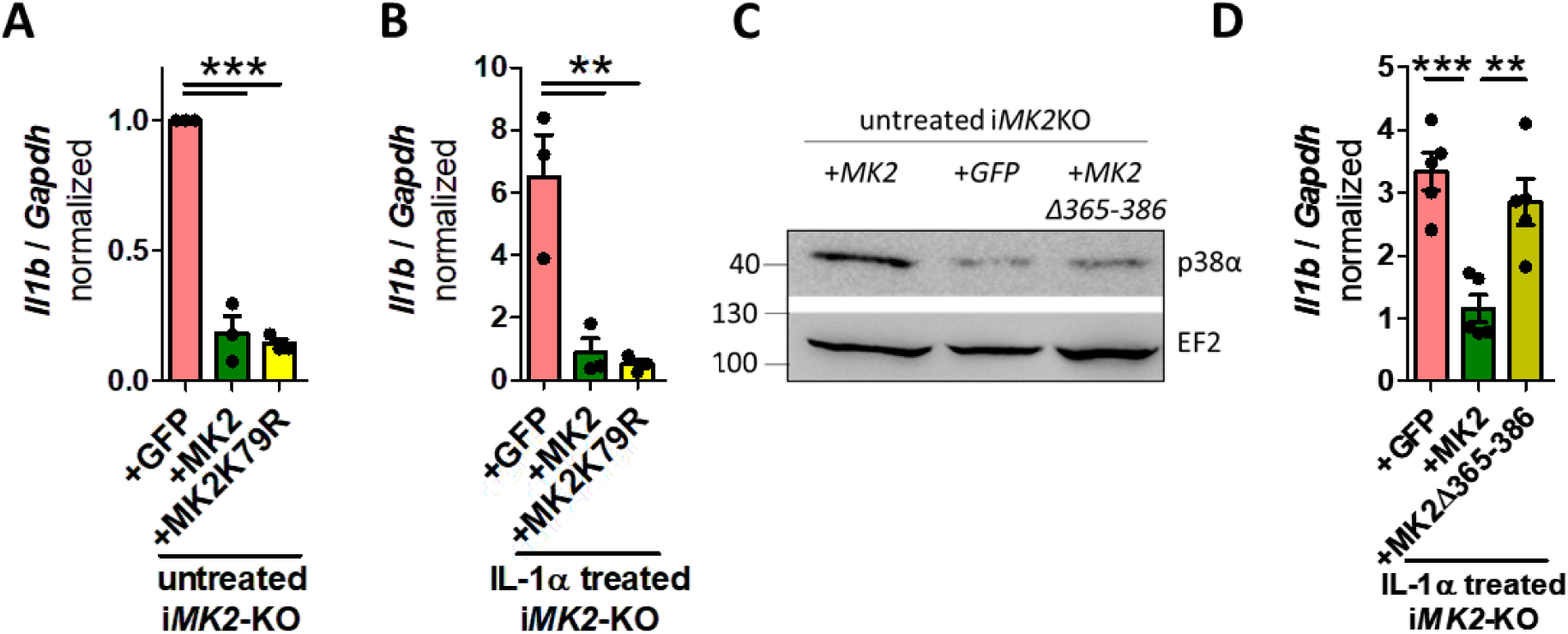
The MK2 kinase activity is not involved, but the MK2 C-terminus is important. **(A)** The rescued MK2 kinase-inactive mutant, MK2K79R, reduces *Il1b* mRNA levels to a degree comparable to that of the rescued MK2 in untreated (UT) and **(B)** IL-1α-treated (5 ng/ml, 1h) i*MK2*-KO macrophages. **(C)** i*MK2*-KO cells have lower levels of the p38α protein. These levels can be restored by rescuing *MK2*, but not by rescuing a MK2 mutant lacking the C-terminus *MK2-Δ365-386*. **(D)** MK2-Δ365-386 does not affect the *Il1b* mRNA levels in IL-1α-treated cells. 1W-ANOVA with Tukey’s Multiple Comparison Test, mean ± SEM, ** P<0.01, *** P<0.001.

### p38α suppresses the non-canonical NF-κB pathway independent of the kinase activity

Overexpression of *p38α* in i*MK2*-KO cells significantly increased basal TRAF2 levels and reduced basal RelB protein levels (Fig. 7A-C). To investigate if the kinase activity of p38α influences the basal non-canonical NF-κB pathway, in addition to *p38α*, a kinase-inactive mutant *p38-AGF* was overexpressed in i*MK2*-KO cells. Surprisingly, basal *Il1b* mRNA was lowered in i*MK2*-KO+*p38α* as well as in i*MK2*-KO+*p38-AGF* cells (Fig. 7D). In addition, *Map3k14* mRNA decreased and *Traf2* mRNA increased (Fig. 7E-F). Also, *Relb, Nfkb2,* and *Il1b* were significantly reduced in IL-1α-treated i*MK2*-KO cells with overexpressed *p38α* and *p38-AGF* (Fig. 7G-H, Suppl. 4F).

**Figure 7:**
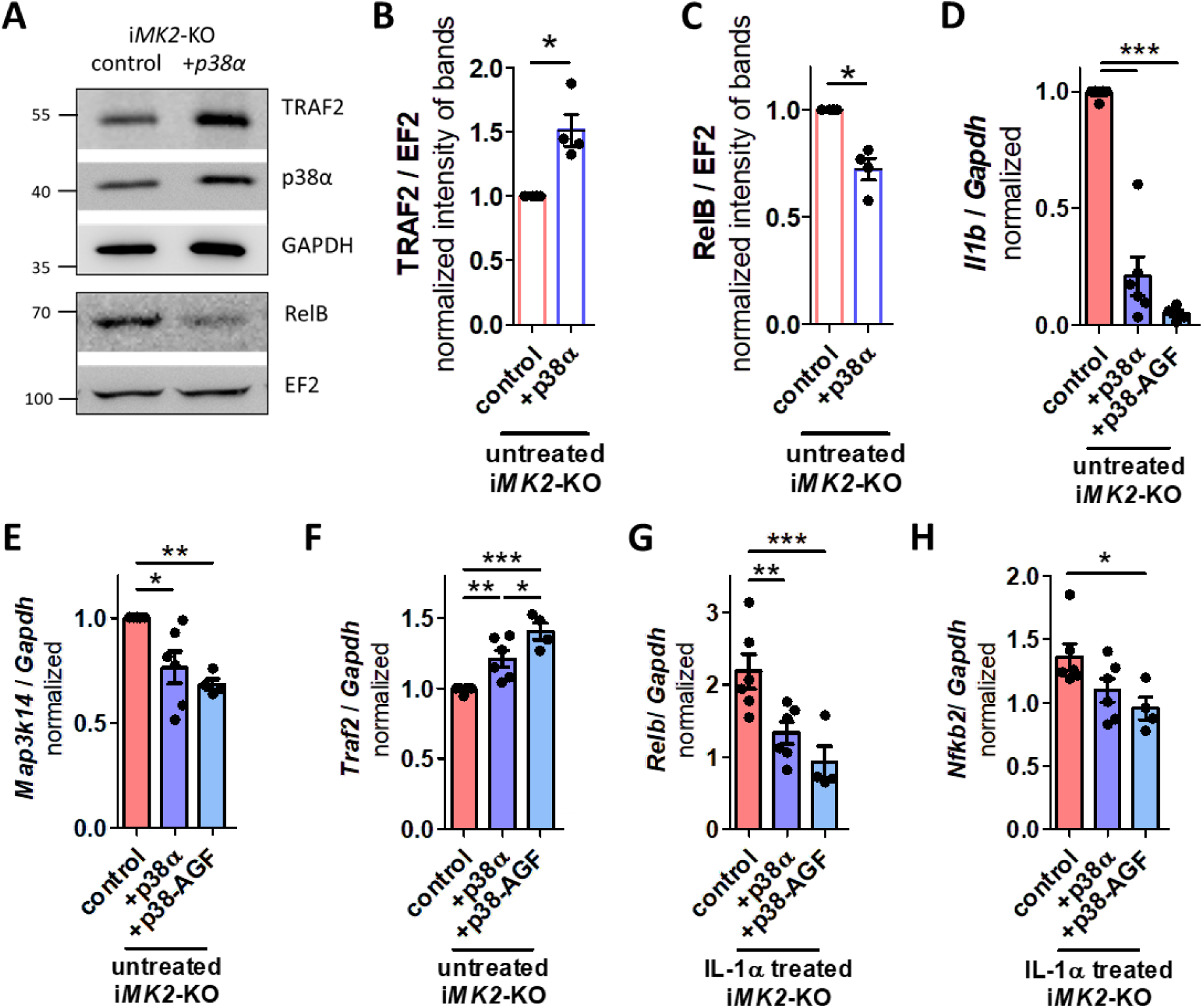
p38α inactivates the non-canonical NF-κB pathway independent of the kinase activity. **(A-C)** Overexpression of *p38α* in i*MK2*-KO cells increases basal TRAF2 and reduces basal RelB protein levels. **(D)** Overexpression of *p38α* and kinase inactive mutant *p38-AGF* reduce basal *Il1b* and **(E)** *Map3k14* mRNA and **(F)** increase basal *Traf2* mRNA in resting i*MK2*-KO cells. **(G)** *Relb* and **(H)** *Nfkb2* mRNA are reduced in IL-1α-treated (5 ng/ml, 1h) i*MK2*-KO+*p38α* and +*p38-AGF* cells. **(B-C)** students t-test, **D-H)** 1W-ANOVA followed by Tukey’s Multiple Comparison Test, mean ± SEM, * P<0.05, ** P<0.01, *** P<0.001.

We then generated two *p38α*-KO RAW 264.7 macrophage cell lines by using CRISPR/Cas9 technology, followed by single-cell sorting. Both *p38α*-KO cell lines showed increased basal mRNA levels of the non-canonical NF-κB signaling components *Map3k14*, *Relb,* and *Nfkb2* (Fig. 8A-C). Additionally, both *p38α*-KO cell lines exhibited significantly higher basal levels of the non-canonical NF-κB signaling proteins NF-κB2 and RelB compared to the control (Fig. 8D-E). In contrast, the protein level of the canonical NF-κB signaling protein RelA did not change in unstimulated *p38α*-KO RAW 264.7 macrophages (Suppl. 4G-H). *p38α*-KO RAW macrophages had a reduced MK2 protein level (Fig. 8D).

**Figure 8:**
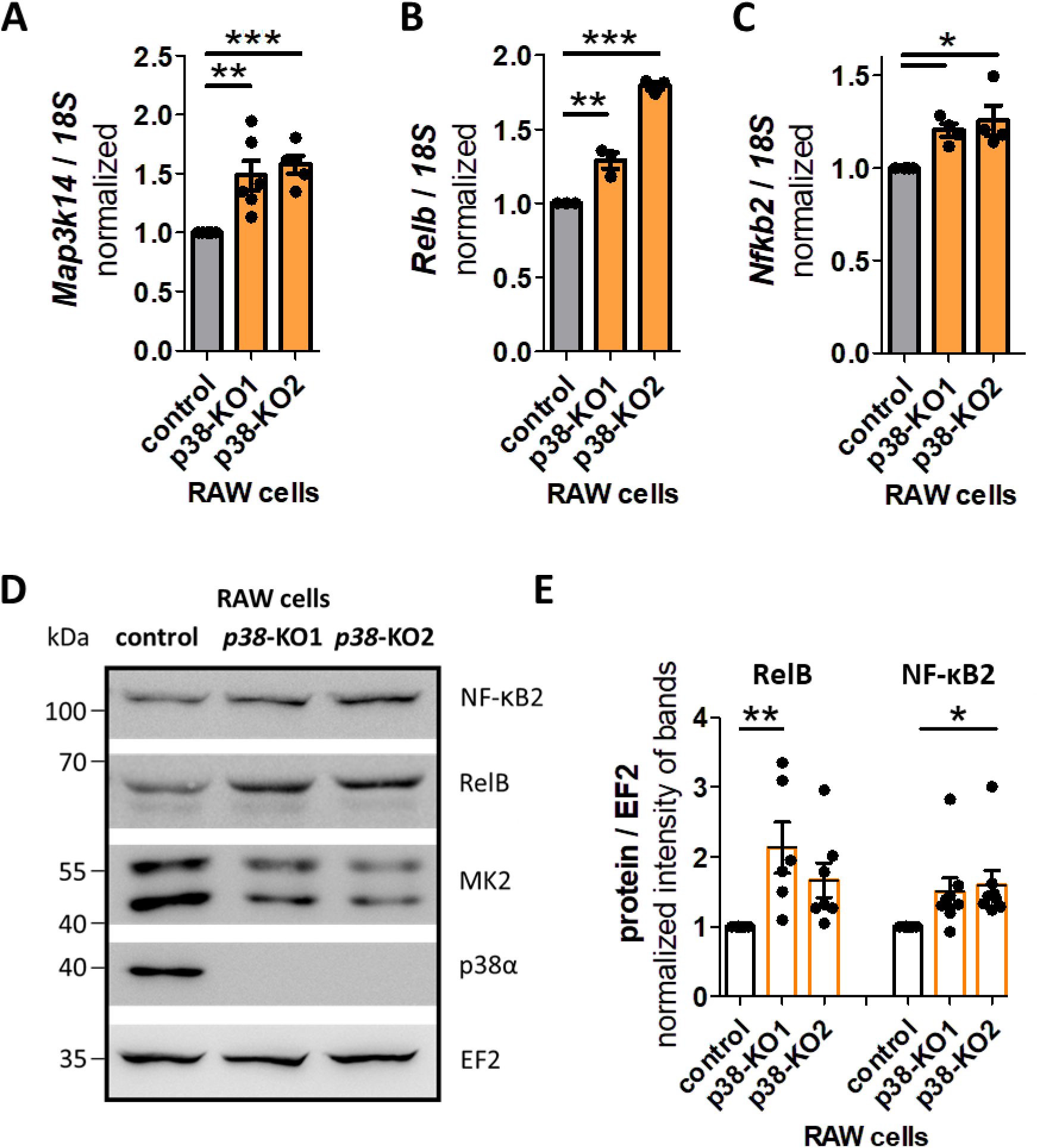
Resting p38α-ko cells have elevated levels of non-canonical NF-κB pathway components. **(A)** Resting *p38α*-KO RAW cells harbor increased *Map3k14* mRNA, **(B)** *Relb* mRNA, and **(C)** *Nfkb2* mRNA levels. **(D-E)** Resting *p38α*-KO RAW cells have elevated RelB and NF-κB2 protein levels. 1W-ANOVA followed by Dunnett’s Multiple Comparison Test, mean ± SEM, * P<0.05, ** P<0.01, *** P<0.001.

Overexpression of p38α and the kinase-inactive *p38-AGF* in i*MK2*-KO cells decreased the levels of components of the non-canonical NF-κB pathway, such as *Map3k14* (NIK), *Relb* and *NFkB2*. Consistent with this finding, knocking out *p38α* in a different macrophage cell line resulted in elevated levels of these components. Therefore, MK2 suppresses the non-canonical NF-κB pathway and resulting IL-1β production independently of its own kinase activity by stabilizing the protein kinase p38α.

### MK2/p38α stabilize p53, which suppresses the non-canonical NF-κB pathway, *Il1b*, and its own expression

p38α seems to regulate the basal non-canonical NF-κB pathway independent of its kinase activity, therefore stabilizing another protein might be important. It has been described, p38 and the tumor suppressor p53 exist within the same physical complex and co-expression of p38 stabilizes p53 protein complex (*12*). In contrast, there are also p38α kinase dependent interactions described. Activation of p38 MAPK has been shown to induce rapid degradation of the E3 ubiquitin-protein ligase mouse double minute 2 homolog (MDM2), thereby stabilizing its target, p53 (*23*). Additionally, p38 has been demonstrated to phosphorylate the tumor suppressor p53 at residues Ser^33^ and Ser^46^ in the N-terminus (*12*).

Therefore, we analyzed the protein levels of p53 and MDM2 in i*MK2*-KO cells. The basal p53 protein level was indeed significantly reduced in i*MK2*-KO cells. However, the basal MDM2 protein level remained unchanged (Fig. 9A-B, Suppl. 4I). We also analyzed the mRNA level of the p53 gene *TP53*. Unlike the reduced p53 protein levels, the *TP53* mRNA levels increased in resting i*MK2*-KO cells (Fig. 9C). This may be due to the previously described increase in RelB protein levels in i*MK2*-KO cells, since *TP53* is a direct target of RelB (*24*). Consistent with this finding, *p38α*-KO macrophages exhibited reduced basal levels of p53 protein, but no significant change in MDM2 protein levels (Fig. 9D-E, Suppl. 4J). Similar to i*MK2*-KO cells, *p38α*-KO macrophages also showed elevated *TP53* transcript levels (Fig. 9F). Additionally, inhibiting p38 kinase activity in resting RAW cells with the specific p38 MAPK inhibitor BIRB 796 (Suppl. 5A) did neither reduce p53 levels nor increase RelB and IL-1β protein levels (Fig. 9G-I, Suppl. 5B-C). Therefore, the basal p53 protein level is stabilized by the MK2-p38-p53 protein interaction, and phosphorylation events are not required.

**Figure 9:**
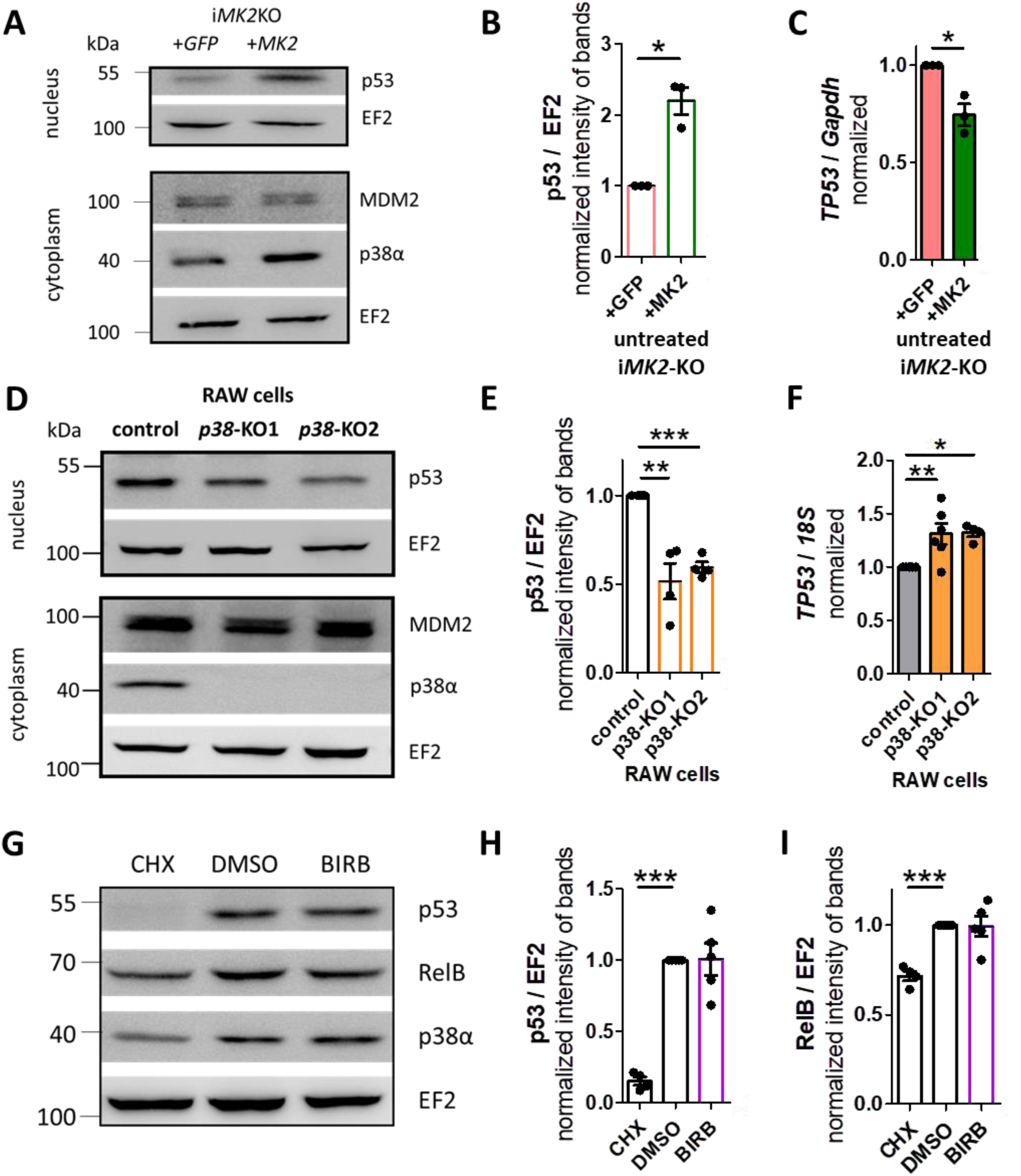
MK2/p38α stabilize p53 protein. **(A-B)** Resting i*MK2*-KO+*GFP* cells have less p53 protein level in the nucleus fraction, but similar MDM2 protein level in the cytoplasmic fraction compared to i*MK2*-KO+*MK2* macrophages. **(C)** *TP53* mRNA is increased in i*MK2*-KO cells. **(D-E)** Resting *p38α*-KO RAW macrophages have lower levels of p53 protein in the nuclear fraction, but similar levels of MDM2 protein in the cytoplasmic fraction compared to the control cells. **(F)** *TP53* mRNA is increased in *p38α*-KO cells. **(G-I)** Effect of the translation inhibitor cycloheximide (CHX, 40 µg/mL, 5h) and the p38 inhibitor BIRB796 (1 µM, 5h) on the levels of the p53 and RelB proteins in the nuclear fraction of RAW264.1 cells. **A-C)** students t-test, **D-I)** 1W-ANOVA followed by Dunnett’s Multiple Comparison Test, mean ± SEM * P<0.05, ** P<0.01, *** P<0.001.

To investigate whether p53 also influences the non-canonical NF-κB pathway, we treated i*MK2*-KO cells with Nutlin-3. Nutlin-3 is an inhibitor that prevents MDM2 from binding to p53, thereby blocking the subsequent degradation of p53 (*25*). As expected, Nutlin-3 treatment resulted in p53 protein accumulation in i*MK2*-KO cells (Fig. 10A). Nutlin-3 also stimulated RelB protein cleavage, as indicated by a weaker RelB band and the appearance of several lower-molecular-weight RelB bands (Fig. 10A-C). Similarly, Nutlin-3-treated RAW 264.7 macrophages also exhibited increased p53 and reduced RelB protein levels (Suppl. 5D-E). No changes in NF-κB2 or TRAF2 protein levels were observed in i*MK2*-KO cells after Nutlin-3 treatment (Fig. 10A). Additionally, *Il1b* and *TP53* mRNA levels were significantly reduced in Nutlin-3-treated i*MK2*-KO cells (Fig. 10D-E). Four hours after Nutlin-3 treatment, *Map3k14*, *Relb,* and *Nfkb2* mRNA levels were also significantly reduced, though *Traf2* levels were not (Suppl. 5F-I). Therefore, p53 suppresses the non-canonical NF-κB pathway by cleaving RelB, thereby inhibiting basal *Il1b* and *TP53* expression.

**Figure 10:**
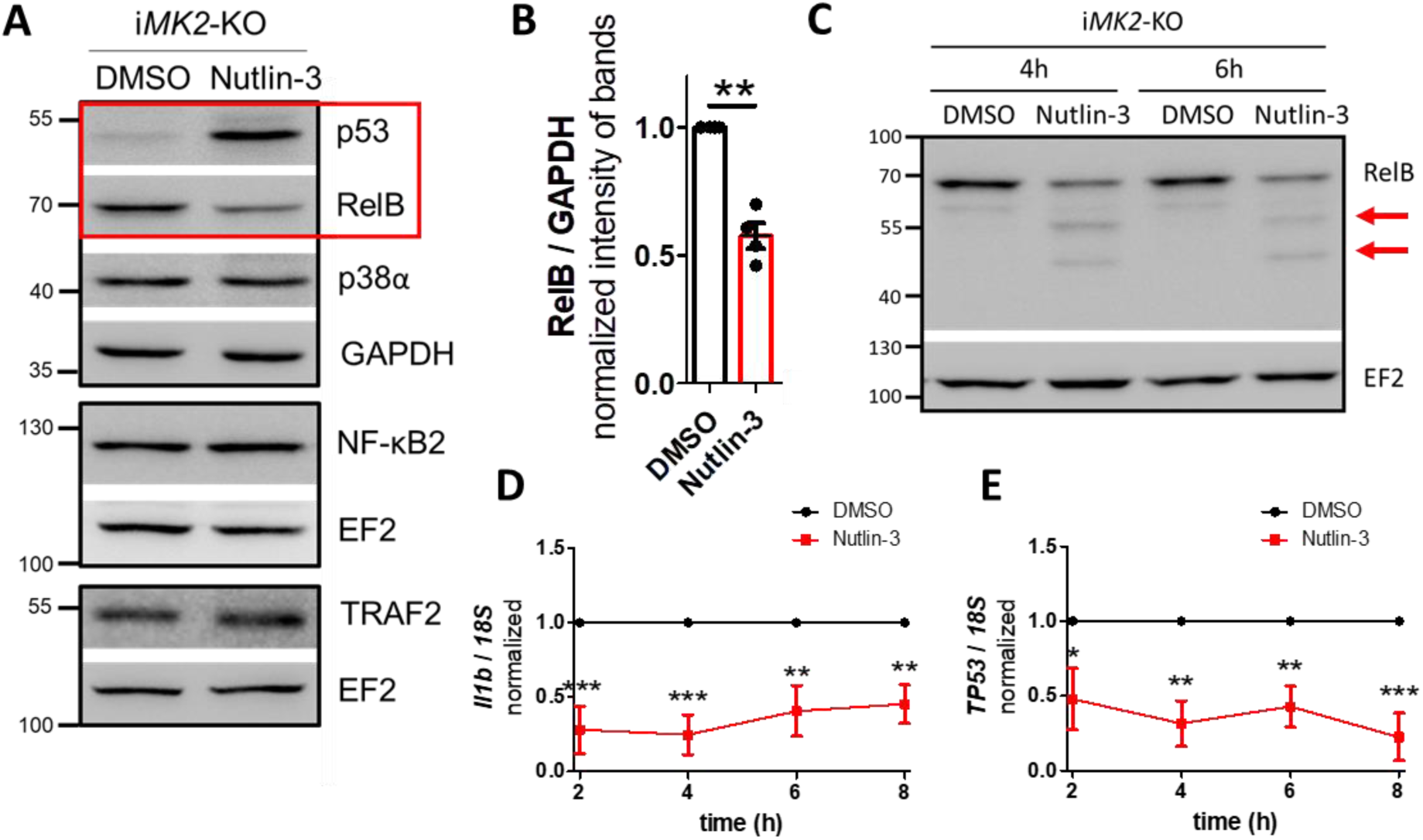
p53 inactivates the non-canonical NF-κB pathway through RelB cleavage. **(A-B)** Nutlin-3 (20 µM, 4h) treated i*MK2*-KO cells accumulate p53 and harbor reduced RelB protein, but not NF-κB2, TRAF2 or p38α. **(C)** RelB cleavage products appear in Western blots after Nutlin-3 (20 µM) treatment in i*MK2*-KO cells. **(D)** *Il1b* mRNA and **(E)** *TP53* mRNA are reduced in Nutlin-3 (20 µM)-treated i*MK2*-KO cells. **B)** students t-test, **D-E)** n=4, 2W-RM-ANOVA with Bonferroni posttests, mean ± SEM, * P<0.05, ** P<0.01, *** P<0.001.

### p53 activates mitochondrial caspase-3, which induces RelB cleavage

Since RelB can be cleaved by caspase-3 (*26, 27*), we analyzed the level of this protease in i*MK2*-KO cells. Following Nutlin-3 treatment, i*MK2*-KO cells exhibited decreased levels of full-length caspase-3 protein (procaspase-3) and increased levels of cleaved Caspase-3 (Fig. 11A-B, Suppl. 5J). Furthermore, the caspase-3-specific inhibitor zDEVDfmk and the pan-caspase inhibitor zVADfmk inhibited Nutlin-3-induced RelB cleavage in i*MK2*-KO macrophages (Fig. 11C-D). Because p53 activates caspases by releasing of mitochondrial cytochrome c (*28*), i*MK2*-KO cells were pre-incubated with either Pifithrin µ (a specific inhibitor of mitochondrial p53 translocation), or Pifithrin α (a p53-specific transcriptional inhibitor). Pifithrin µ, but not Pifithrin α, reduced Nutlin-3-induced RelB cleavage (Fig. 11E-F). Therefore, p53 activates caspase-3 through the mitochondrial pathway, which cleaves the non-canonical RelB protein. To confirm this non-transcriptional effect, a prominent transcriptional target of p53, the cyclin-dependent kinase inhibitor p21, was analyzed. There was no significant difference in the protein levels of p21 in the nuclear fraction of resting *p38*-KO and control cells (Suppl. 5K-L).

**Figure 11:**
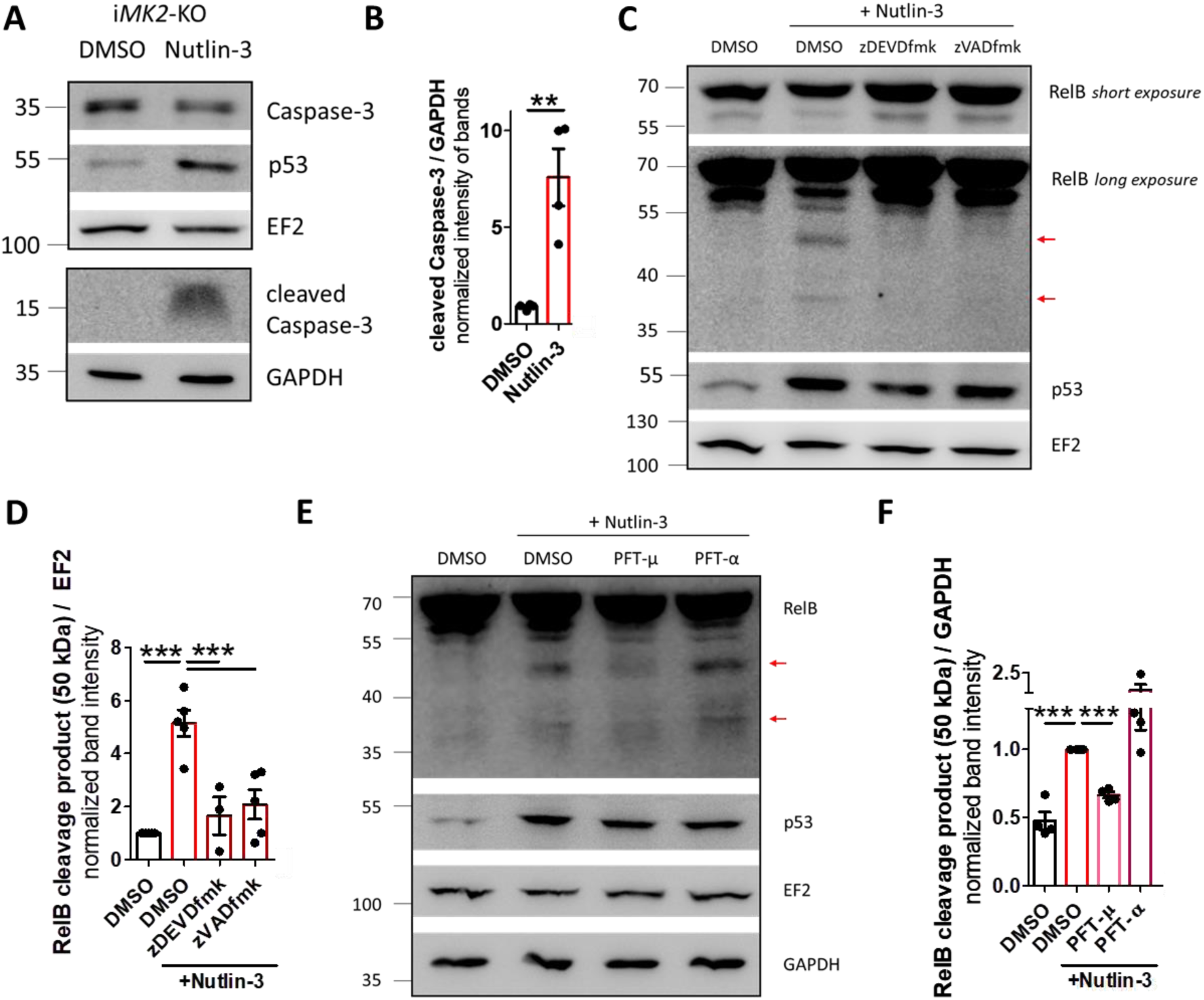
p53 activates mitochondrial caspase-3, which cleaves RelB. **(A-B)** Nutlin-3 (20 µM, 4h) treated i*MK2*-KO cells have reduced levels of full-length Pro-caspase-3 and increased levels of cleaved Caspase-3 protein. **(C-D)** The appearance of RelB cleavage products after Nutlin-3 (20 µM, 4h) treatment in i*MK2*-KO cells can be blocked by the caspase-3 inhibitor z-DEVD-fmk (100 µM, 1h), the pan-caspase inhibitor z-VAD-fmk (25 µM, 1h), and **(E-F)** the inhibitor of p53 mitochondrial translocation Pifithrin-µ (10 µM, 1h), but not by the p53 transcriptional inhibitor Pifithrin-α (10 µM, 1h) in i*MK2*-KO cells. **(B)** students t-test, **(D-F)** 1W-ANOVA followed by Dunnett’s Multiple Comparison Test, mean ± SEM, ** P<0.01, *** P<0.001.

Since treatment with Nutlin-3 also leads to RelB cleavage in WT RAW 264.7 macrophages (Suppl. 5D-E), MK2 and p38α contribute to RelB cleavage exclusively through the stabilization of the p53 protein. In presence of MK2 and p38, p53 is stabilized within the MK2-p38-p53 complex. Stabilization of p53 activates mitochondrial caspase-3, which cleaves RelB and keeps therefore the activity of the non-canonical NF-κB pathway diminished. In contrast, the absence of MK2 or p38α destabilizes the p53 protein. This leads to an increased RelB protein level and activation of the non-canonical NF-κB pathway in resting *MK2*-KO and *p38*-KO cells (Fig. 12).

**Figure 12:**
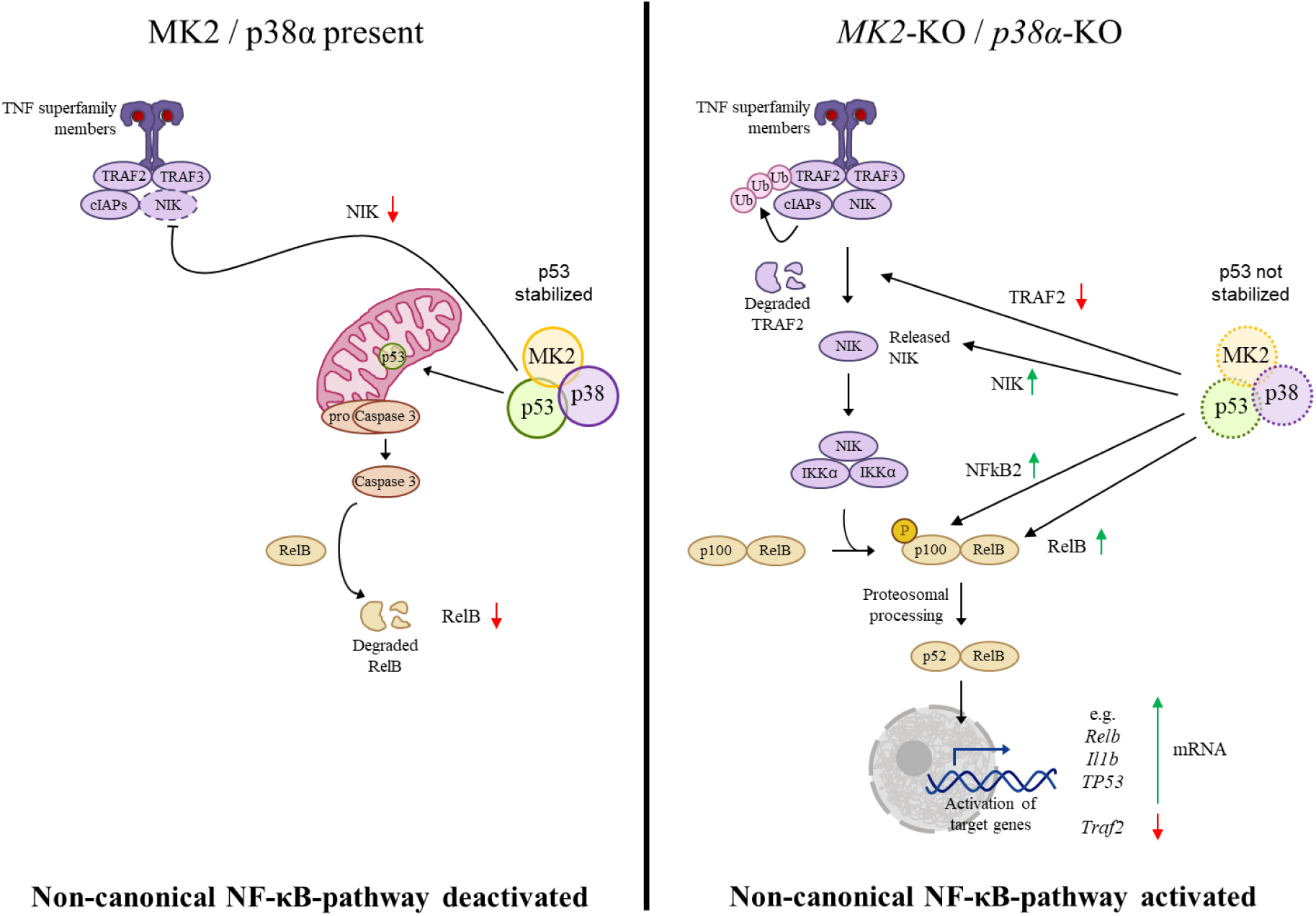
Synopsis. The tumor suppressor p53 is known to negatively regulate the non-canonical NF-κB pathway by suppressing the expression of *Map3k14* (NIK) mRNA. This study identifies a novel mechanism by which p53 inhibits the non-canonical NF-κB pathway in macrophages: p53 activates caspase-3, which subsequently cleaves RelB. Absence of the protein kinase complex MK2/p38α destabilizes p53, leading to the basal activation of the non-canonical NF-κB pathway.

- The protein kinases MK2 and p38α stabilize the level of the p53 protein in macrophages.
- p53 activates mitochondrial caspase-3, which cleaves the non-canonical NF-κB component RelB.
- Resting *MK2*-KO and *p38α*-KO macrophages display reduced basal levels of p53 and increased activation of the non-canonical NF-κB pathway.
- In resting *MK2*-KO cells, the non-canonical NF-κB pathway suppresses TRAF2 expression and induces RelB transcription, as well as that of the cytokine IL-1β and p53 itself.

## Discussion

IL-1β plays an important role in maintaining human homeostasis by regulating nutrition, sleep, and temperature (*29*). However, it is also a potent endogenous pro-inflammatory pyrogen that influences the innate and adaptive immune systems. Prolonged, uncontrolled release of IL-1β causes the pathological effects of chronic inflammation. It is involved in the pathogenesis of various diseases, including arthritis, gout, Alzheimer’s disease, diabetes mellitus, obesity, atherosclerosis, and tumor growth (*30–33*). The expression of the inactive pro-IL-1β precursor is usually triggered when pattern recognition receptors, such as Toll-like or NOD-like receptors, are stimulated by microbial or viral molecules (*34, 35*). However, IL-1β can also stimulate its own expression. The main subunits responsible for activating the NF-κB binding site in the *Il1b* promoter region in response to phorbol 12-myristate 13-acetate (PMA) or LPS stimulation are key components of the canonical NF-κB pathway, including NF-κB1 (p50), RelA (p65), and c-Rel (*36–38*). Nevertheless, RelB competes with RelA for binding to κB sites of NF-κB-regulated promoters in dendritic cells (*39*). Additionally, increased *Il1b* levels are detected in *Relb*-KO fibroblasts (*40*). In both cases, RelB suppresses LPS-induced inflammatory gene expression. Accordingly, RelB has been shown to prevent RelA from binding to the *Il1b* promoter in LPS-tolerized cells by forming facultative heterochromatin, which silences *Il1b* transcription (*41–43*).

In this study, we demonstrate the activation of *Il1b* transcription through the non-canonical NF-κB pathway in resting i*MK2*-KO macrophages. The abundance of the non-canonical NF-κB pathway is enhanced in these cells, since basal TRAF2 expression is reduced while basal *Map3k14* (NIK), RelB and NF-κB2 expressions are increased. However, components of the canonical pathway, such as RelA and NF-κB1, remain unaffected. Additionally, mRNA analysis of unstimulated i*MK2*-KO macrophages revealed elevated levels of the targets of the non-canonical NF-κB pathway, including *Il1b*. Inhibiting the non-canonical NF-κB pathway significantly reduced *Il1b* in resting i*MK2*-KO cells, but not in LPS-stimulated cells. In contrast, inhibiting the canonical NF-κB pathway did not lower the basal *Il1b* level in i*MK2*-KO cells, but LPS-induced *Il1b* levels were reduced. These results confirm that LPS activates the canonical NF-κB pathway and *Il1b* transcription. However, the results also reveal an as-yet-unknown activation of *Il1b* transcription through the non-canonical NF-κB pathway in quiescent i*MK2*-KO cells. Similar activation of the *Il1b* promoter, independent of the canonical NF-κB pathway, has only been described for STAT3 so far (*44*). Additionally, STAT3 has been shown to activate the non-canonical NF-κB pathway (*45–47*). Furthermore, 2,3,7,8-tetrachlorodibenzo-p-dioxin (TCDD) has been shown to induce *Il1b* promoter activity in a RelB-dependent manner. However, the underlying molecular mechanisms remain unclear (*48*).

Targeting the human MK2-p38 complex with Zunsemetinib (ATI-450) has been attempted to inhibit IL-1β (*49*). However, at low concentrations where ATI-450 specifically inhibits the MK2-p38 complex, IL-1β release from human peripheral blood mononuclear cells (PBMCs) significantly increases. The study above does not discuss the increase in IL-1β levels at low ATI-450 concentrations, but these data align with our findings on the activation of the non-canonical NF-κB pathway and subsequent increase in basal IL-1β secretion. This may also explain why a recent phase IIb trial of ATI-450 in rheumatoid arthritis did not meet the expected endpoints.

We demonstrate that the activation of the non-canonical NF-κB pathway in resting i*MK2*-KO cells is caused by the destabilization of the p38 and p53 proteins. Accordingly, increased basal levels of non-canonical *Map3k14* (NIK), RelB, and NF-κB2 were detected in *p38α*-KO RAW cells. Levels of the p53 protein were significantly reduced in both, immortalized macrophages derived from *MK2*-KO mice, as well as in CRISPR-edited *p38α*-KO RAW 264.7 cells. MK2 is known to phosphorylate MDM2, leading to reduced p53 protein levels. In *MK2*-KO fibroblasts, p53 protein levels increase following exposure to UV radiation or treatment with anisomycin (*50*). These stimuli activate various protein kinases besides p38 and MK2 (*51*). Another study found that skin biopsies of *MK2*-KO mice revealed increased p53 protein levels, as well as reduced *Il1b* mRNA and IL-1β protein levels after treatment with 7,12-dimethylbenz[a]anthracene (DMBA)/12-O-tetradecanoylphorbol-13-acetate (TPA). Additionally, MDM2 phosphorylation was reduced in *MK2*-KO keratinocytes in vitro (*52*). While these studies analyzed different cell types (such as fibroblasts and keratinocytes) and used different stimuli (such as anisomycin or UV radiation), we examined basal levels of IL-1β and p53 in unstimulated *MK2*- or *p38α*-KO macrophages. This explains the opposing effects induced by protein-protein stabilization versus effects dependent on the kinase activity of MK2/p38. Indeed, p38 and p53 are located in the same physical complex, and co-expression of p38 stabilizes p53 (*53*). On the other hand, p53-stabilizing effects that depend on p38 kinase activity have also been described. In a cell line that is resistant to the chemotherapeutic agent paclitaxel, constitutively activated p38 induces the rapid degradation of MDM2, thereby stabilizing p53 (*23*). Additionally, p38 phosphorylates p53 at Ser^33^ and Ser^46^ following UV exposure, which may contribute to p53 stabilization (*53*). In conclusion, opposing effects in the stabilization of p53 by MK2 or p38 have been described, highlighting the complexity and cell-type specificity of p53 regulation.

Adding Nutlin-3 to cells inhibits MDM2 and the subsequent degradation of p53. Nutlin-3 treatment also inhibits the non-canonical NF-κB pathway by cleaving RelB in i*MK2*-KO cells and RAW macrophages. RelB is a direct transcription factor of *TP53* (*24*). Indeed, Nutlin-3 treatment also reduced *TP53* mRNA and *Il1b* mRNA. The tumor protein p53 activates caspase-9/-3 through the release of mitochondrial cytochrome c (*54–56*). In addition, activated caspase-3 can cleave RelB (*27, 57*). Indeed, treatment with Nutlin-3 activates caspase-3, as evidenced by decreased levels of full-length pro-caspase 3 and increased levels of cleaved caspase-3. Inhibiting the p53 mitochondrial pathway or caspase-3 itself prevents RelB cleavage following Nutlin-3 treatment.

After activation, the non-canonical NF-κB pathway has been shown to regulate the expression of its own components. RelB can bind to its own promoter NF-κB sites, thereby autoregulating *Relb* mRNA levels (*58*). Furthermore, RelB has been shown to repress *Traf2* expression (*59*). This repression prevents subsequent NIK degradation, which in turn activates the non-canonical NF-κB pathway. Decreased levels of the p53 protein and activation of caspase-3 were observed in *Traf2*-deficient cells due to activated JNK signaling (*60*). In addition to RelB, NF-κB2 is also positively autoregulated (*61*). To our knowledge, no study has yet been published regarding the positive regulation of RelB or NF-κB2 with respect to their binding to the NIK promoter region or *Map3k14* (NIK) expression. This study describes the significantly reduced *Map3k14* mRNA expression in i*MK2*-KO cells after inhibition of the non-canonical NF-κB pathway. This finding suggests that the non-canonical NF-κB pathway indirectly regulates *Map3k14* mRNA levels. It was shown that p53 suppresses the mRNA expression of *Map3k14* through the miRNA pathway. Knockdown of the tumor suppressor p53 stabilizes NIK, whereas accumulation of p53 protein decreases endogenous NIK protein (*62*). The regulation of NIK transcription by p53 could explain our results, as we observed decreased basal p53 protein levels and increased basal *Map3k14* mRNA expression in i*MK2*-KO and *p38α*-KO macrophages. Additionally, we measured significantly reduced *Map3k14* mRNA levels in Nutlin-3-treated RAW cells with p53 accumulation.

Here we show, that p53 negatively regulates the non-canonical NF-κB pathway through caspase-3-dependent cleavage of RelB. In i*MK2*-KO and *p38α*-KO macrophages, accompanied by reduced p53 protein levels, the non-canonical NF-κB pathway is activated. This pathway increases the expression of its target genes, including components of the non-canonical NF-κB pathway itself, as well as *Il1b* and *TP53*. Overexpression of *p38α* in i*MK2*-KO cells rescued the protein levels of the non-canonical NF-κB pathway components TRAF2 and RelB. Treatment with Nutlin-3 induced p53 protein accumulation and blocked the non-canonical NF-κB pathway by cleaving RelB with caspase-3. Subsequently, the expression of the targets of the non-canonical NF-κB pathway as well as *Il1b* and *TP53* mRNA, decreased. Thus, p53 can downregulate its own *TP53* expression. This reveals a new autoregulatory mechanism of p53 protein and *TP53* mRNA levels through RelB cleavage. These results uncover a new mechanism contributing to the long-discussed link between cancer and inflammation, in which the tumor suppressor p53 represses cytokine expression. Consequently, the loss of p53 would lead to oncogenic transformation, increased basal cytokine levels and chronic inflammation confirming Virchow’s classical observation that inflammation and neoplasia occur coincidentally.

## Materials and Methods

### Cell lines and growth conditions

The wild-type C57BL/6, MK2-, and MK2/3-knockout strains (*63, 64*) were maintained under specific pathogen-free conditions at the Hannover Medical School animal facility. All animal experiments were approved by the appropriate institutional and state animal welfare committees. Procedures were conducted in accordance with local animal use and care committee guidelines, as well as national animal welfare laws. Both male and female adult mice (ages 7 weeks to 9 months) were used. Blood was collected postmortem from the vena cava and heart. After 30 min of clotting, the serum was collected by centrifugation (1500xg, 10 min, 4°C) and stored at -20°C. Bone marrow cells were isolated and differentiated into bone marrow-derived macrophages (BMDMs) in DMEM (Gibco) supplemented with 10% fetal bovine serum (FBS, Capricorn Scientific), penicillin/streptomycin (Capricorn Scientific), Gentamycin (10 μg/ml) / Amphotericin B (0,25 μg/ml) solution (CELLnTEC), MEM non-essential amino acids (Gibco), and rM-CSF (80 ng/ml, Wyeth, Boston, MA) for 8 days in a humidified incubator at 37°C and 5% CO₂. The cells were harvested by incubating them for 30 min with non-enzymatic cell dissociation solution (C5789, Sigma-Aldrich), followed by scraping, washing and counting. Experiments were performed the next day. The N-values of BMDMs of the same genotype differ in certain experiments because there were not enough cells available for some mice to perform all assays in parallel, and some experiments were performed at a different time with other mice of the same genotype.

*p38α*-KO cells were generated by transfecting RAW 264.7 cells (*M. musculus*, CVCL_0493) with either the *p38α MAPK14* Double Nickase Plasmid (sc-424051-NIC) or the Control Double Nickase Plasmid (sc-437281), according to the Santa Cruz Biotechnology (SCBT) protocol. Then, the cells were sorted by single-cell sorting of GFP-positive cells. To generate MK2 knockdown, RAW cells were treated with either control siRNA (sc-37007) or MK2 siRNA (sc-35856 according to SCBT protocol.

Immortalized MK2-KO (iMK2-KO) macrophages (*M. musculus*) were rescued through viral transduction with an empty vector (pMMP-IRES-GFP), MK3, MK2, a kinase-inactive MK2 mutant (MK2K79R) (*5, 65*), or a mutant lacking an NcoI recognition site and the MK2 C-terminus pMMP-IRES*-MK2-K357R-Δ365-386* (termed as *MK2-Δ365-386)*. The cells were then sorted by the GFP-positive signal. p38α and *p38-AGF* overexpression in iMK2-KO cells was achieved through viral transduction using either an empty pMSC vector (control), or a pMSC vector containing p38α. The cells were selected using 100 μg/mL of hygromycin (Roth) for 9 days. Amphotropic retroviruses were generated by transfecting the 293 GPG packaging cell line (*Homo sapiens*, *66*) with the indicated vectors. The RAW 264.7 and iMK2-KO cells were cultured in DMEM supplemented with 10% FBS and penicillin/streptomycin.

### RNA purification and analysis

BMDM (5x10^5^ cells/well), i*MK2*-KO or RAW 264.7 cells (both 2x10^5^ cells/well) were seeded, and treated one day later with the indicated concentrations and durations of recombinant murine IL-1α (211-11A, Peprotech), LPS (Escherichia coli 0127:B8; Sigma-Aldrich), Actinomycin D (Cayman Chemicals), B022 (MedChemExpress), BMS 345541 (Axon Medchem), CAY10657 (Cayman Chemicals), SML1160 (Sigma-Aldrich), IKK inhibitor XII (HPN-01, Sigma Aldrich), sc-514 (Cayman Chemicals), T-5224 (Cayman Chemicals), Takinib (MedChemExpress) or Nutlin-3 (SCBT).

RNA was extracted using TRIzol Reagent (Invitrogen) and 0.5 µg RNA (according to the NanoDrop ND-1000 spectrophotometer, peqlab Biotechnologie) was converted to cDNA through reverse transcriptase PCR with Biometra TRIO (analytik jena), contained RevertAid Reverse Transcriptase (EP0442), Oligo(dT)18 Primer (SO132), RiboLock RNase Inhibitor (EO0381) (all from Thermo Fisher Scientific), and Deoxynucleotide (dNTP) Solution Mix (N0447S, New England Biolabs). The cDNA was diluted 1:10, and quantitative PCR was performed using a Rotor-Gene Q (Quiagen), qPCRBIO SyGreen® Mix Separate-ROX (PCR Biosystems), along with the following mouse primers: Gapdh fw 5’-CAT GGC CTT CCG TGT TCC TA-3’, rev 5’-CCT GCT TCA CCA CCT TCT TGA T-3’; Il1b fw 5’-GTG ATA TTC TCC ATG AGC TTT G-3’, rev 5’-TCT TCT TTG GGT ATT GCT TG-3’; Tnf fw 5’-CAT CTT CTC AAA ATT CGA GTG ACA A-3’, rev 5’-TGG GAG TAG ACA AGG TAC AAC CC-3’; Relb fw 5’-TAC GAC AAG AAG TCC ACC-3’, rev 5’-TTT TTG CAC CTT GTC ACA G-3’; Nfkb2 fw 5’-CTT CAG ATT TCG ATA TGG CTG-3’, rev 5’-AGT TAC AGA TCT TGA CAG TAG G-3’; Rela fw 5’-GTA TCC ATA GCT TCC AGA AC-3’, rev 5’-GAA AGG GGT TAT TGT TGG TC-3’; Nfkb1 fw 5’-TGT ATT AGG GGC TAT AAT CCT G-3’, rev 5’-ATG ATC TCC TTC TCT CTG TC-3’; Traf2 fw 5’-GAC CTT GAA AGA ATA CGA GAG-3’, rev 5’CAG ACC TCA TAG TGT ACC TC-3’; Traf3 fw 5’-GAC TCT TCT AAG GAG TGA GG-3’, rev 5’-TGG ATG CTC TTG TTT TTC TC-3’; TP53 fw 5’-TAG GTA GCG ACT ACA GTT AG-3’, rev 5’-GGA TAT CTT CTG GAG GAA GTA G-3’; Il1r1 fw 5’-ACA ATT GTA TGC TGT GTG TG-3’, rev 5’-CGT ATG TCT TTC CAT CTG AAG; il1r2 fw 5’-ACC AGT ACT CAG AGA ATG ATG-3’, rev 5’-AAG ATG ATC AGA GAC AGA GG-3’; Il1rn fw 5’-CAG AAG ACC TTT TAC CTG AG-3’, rev 5’-GGC ACC ATG TCT ATC TTT TC-3’; 18S fw 5’-GTA ACC CGT TGA ACC CCA TT-3’, rev 5’-CCA TCC AAT CGG TAG TAG CG-3’; Map3k14 fw 5’-GCT TAC TGA GAA ACT CAA GC -3’, rev 5’-CTA GTT CCT CTA CCC GAA AC -3’; MAPKAPK2 fw 5’CCC AGT TCC ACG TCA AGT CGG GCC TGC-3’, rev 5’-GGT TCT CAG GCT TGA CAT CCC GGT GAG C-3; MAPK14 fw 5’GAC CTT CTC ATA GAT GAG TGG AAG A-3’, rev 5’-CAG GAC TCC ATT TCT TCT TGG T-3’. The primers were purchased from either Eurofins, MWG Biotech AG, or Sigma-Aldrich. The results were obtained using the ΔCT method with normalized threshold values relative to *Gapdh* or *18S*. Significances were calculated and graphs were generated using GraphPad Prism.

Additionally, the Research Core Unit Genomics at Hannover Medical School performed 1x 75 bp single-read RNA sequencing of total RNA of each one sample after ribo-depletion using an Illumina NextSeq 550. Heatmaps were generated using Qlucore Omics Explorer 3.9.

### Protein analysis

BMDM (5x10^5^ cells/well), i*MK2*-KO (6x10^5^ cells/well), or RAW 264.7 cells (1x10^5^ cells/well) were seeded for whole-cell lysis. The next day, the cells were treated with IL-1α, LPS, B022, BIRB 796 (axon medchem), Nutlin-3 (sc-45061, SCBT), zDEVDfmk (HY-12466, Hycultec), zVADfmk (4026865, Bachem), Pifithrin µ (HY-10940, Hycultec), or Pifithrin α (506132, Sigma-Aldrich) as indicated, and then lysed using kinase lysis buffer supplemented with protease inhibitors (B14001, Bimake) and phosphatase inhibitors (B15001, Bimake).

To isolate nuclear and cytoplasmic proteins, rescued i*MK2*-KO (2x10^6^ cells/plate) were seeded and stimulated the next day as indicated. Cyclohexamide (40 µg/ml; Sigma Aldrich), DMSO (Chemsolute) or BIRB 796 were incubated for 5h prior to nuclear protein extraction. NE-PE Nuclear and Cytoplasmic Extraction Reagents (78835, Thermo Scientific) were used for protein isolation and the Pierce BCA Protein Assay Kit (23227, Thermo Scientific) was used to determine protein concentration according to the provided protocols.

20 – 40 µg of protein were mixed with 4× Laemmli’s SDS sample buffer, heated at 95°C for 5 min, separated by SDS-PAGE on 7.5% to 16% gradient gels with a PageRuler Prestained Protein Ladder (26616, Thermo Fisher Scientific) and transferred to Hybond ECL nitrocellulose membranes (GE Healthcare) through semidry blotting. The primary antibodies GAPDH (MAB374, Merck Millipore), IκBa (9242, Cell Signaling Technology (CST)), MDM2 (D-7) (sc-13161, SCBT), EF2 (C-9) (sc-166415, SCBT), NF-κB1 p105/p50 (D7H5M) (12540, CST), NF-κB2 p100/p52 (4882, CST), NF-κB p65 (L8F6) (6956, CST), RelB (C1E4) (4922, CST), p53 (1C12) (2524, CST), c-Rel (D4Y6M) (12707, CST), Mouse IL-1β /IL-1F2 (AF-401-SP, R&D systems), Histone H3 (9715, CST), NIK (4994, CST), TRAF2 (C192) (4724, CST), TRAF-3 Isoform 2 (MAB3278, R&D Systems), MAPKAPK-2 (D1E11) (12155, CST), MAPKAPK-3 (3043, CST), caspase-3 (H-277) (sc-7148, SCBT), cleaved Caspase-3 (Asp175) (9661, CST), p21 Waf1/Cip1 (E2R7A) (37543, CST), Human/Mouse cIAP Pan-specific Antibody (MAB3400, R&D Systems), or p38α Antibody (C-20) (sc-535, SCBT) were incubated overnight at 4°C, followed by a 2h incubation with secondary horseradish peroxidase-conjugated antibodies (SCBT) at room temperature. The chemiluminescence (ECL) detection solution (solution A, 1.2 mM luminol in 0.1 M Tris-HCl [pH 8.6]; solution B, 6.7 mM p-coumaric acid in dimethyl sulfoxide; 35% H_2_O_2_ solution; ratio, 3333:333:1) and the LAS-3000 luminescent image analyzer (Fujifilm) with the associated Image Reader LAS-3000 software were used for detection. The bands were analyzed using Fiji software. Significances were calculated and graphs were generated using GraphPad Prism. Microsoft PowerPoint was used to crop and arrange the Western blots, and to draw the overview image. Western blots were excluded from analysis if the lanes differed in Ponceau staining, if the band intensity of the control proteins (used for normalization) differed, if the bands were too weak, or if the background was too high for analysis with Fiji software. The N values of the Western blots also differ, because some nuclear extracts had too low protein concentration to perform multiple Western blots from the same sample.

### Cytokine measurements and cell viability

BMDMs (1x10^^5^ cells/well) were stimulated with IL-1α, LPS, ATP (tlrl-atpl, Invivogen), or Nigericin (11437, Cayman Chemicals) as indicated. The Mouse IL-1β uncoated ELISA Kit (88-7013-22, Invitrogen) was used to determine the IL-1β concentration in the supernatant. Absorbance was measured using a Cytation 1 instrument (BioTek). Serum of wild-type or *MK2/3* double knockout mice from the same breeding were analyzed for basal IL-1β, TNF-α and CXCL1 using the Fireplex-96 Inflammation (Mouse) Immunoassay panel (ab235659, Abcam), and detected using a BD Accuri C6 flow cytometer. The data were analyzed using BD Accuri C6 and GraphPad Prism software. CellTiter-Blue® Cell Viability Assay (G8080, Promega) was used to determine the cell viability of resting BMDMs. As negative control, cells were treated with 0.1% Triton X-100 (Sigma Aldrich). Absorbance was measured using a Cytation 1 instrument (BioTek).

### Quantification and statistical analysis

The number of animals used and the number of times the experiments were repeated are indicated in the figure legends or shown through dot plots. Representative Western blots are depicted. Biological replicates, not technical replicates, are shown. All experiments were non-randomized. The sample size was not determined in advance. The investigators were not blinded. Statistical analyses were performed using Prism, with either a two-sided paired or unpaired t-test, or two-way repeated measures (2W-RM) ANOVA with Bonferroni post-tests. An F-test was used to calculate variance. Samples were excluded from the analysis, if the housekeeping gene (qRNA) or the control protein (Western blot) was a significant outlier, as determined using the GraphPad outlier calculator. The statistical details for each experiment, including the tests performed and the significance values (* p < 0.05; ** p < 0.01; *** p < 0.001) are provided in the figure legends and figures. Error bars represent the standard error of the mean (SEM). DeepL was used to improve the style and quality of the texts created by the authors. The abstract was shortened using ChatGPT.

## Acknowledgements

The authors thank the Research Core Unit Genomics at Hannover Medical School for providing sequencing services.

## Author contributions

S.M.H., D.S., S.P., L.A.H., A.D., R.N., N.R., T.Y. performed experiments and analyzed the data, S.M.H, A.K. and M.G. designed the project, S.M.H. and M.G. acquired funding and wrote the manuscript.

## Data and materials availability

The RNA-seq data generated in this study are available in the Gene Expression Omnibus (GEO) database under the accession code GSE299040. Any unique materials generated in this study can be requested from the corresponding authors.

## Supplement

Figs. S1 to S5

