## Supplemental figures for "MK2/p38/p53 suppress basal IL-1β and non-canonical NF-κB signaling"

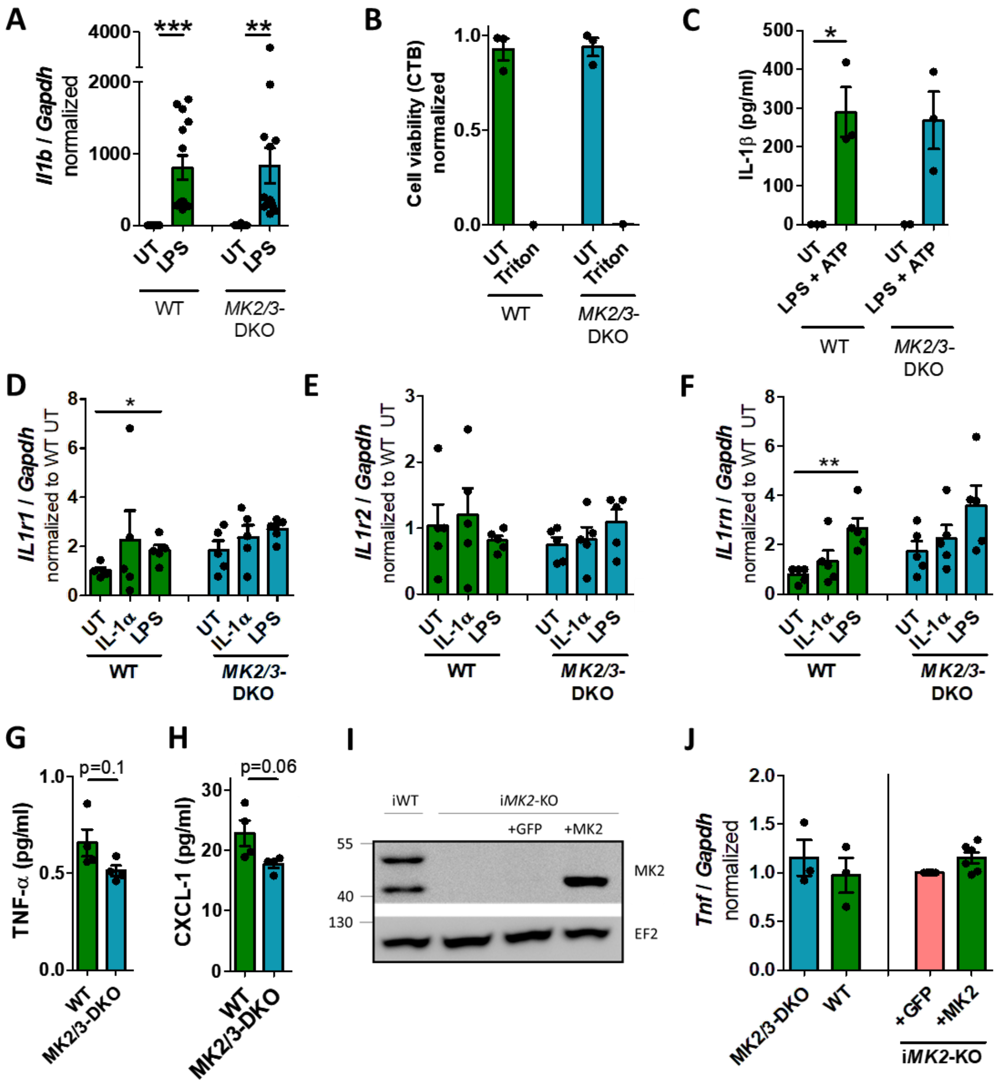


**Supplement Figure 1:**

**(A)** LPS-treated (100 ng/ml, 1h) wild type (WT) and *MK2/3* double-knockout (DKO) bone marrow-derived macrophages (BMDMs) have comparable *Il1b* mRNA levels. WT n=14, DKO n=13. **(B)** Normalized cell viability of resting WT and *MK2/3*-DKO BMDMs was measured using a CellTiter-Blue (CTB) assay. **(C)** The IL-1β concentration in the supernatant of LPS-treated (1 µg/ml, 2h) *MK2/3*-DKO BMDM is similar to that of WT BMDM after the addition of ATP (3 mM, 8h). The mRNA levels of the IL-1 receptors **(D)** *IL1r1*, **(E)** *IL1r2*, and **(F)** *IL1rn* are similar in WT and *MK2/3*-DKO BMDMs. The cells are either untreated (UT), IL-1α (5 ng/ml, 1h) or LPS (100 ng/ml, 1h)-treated. Basal levels of **(G)** Tumor necrosis factor (TNF)-α and **(H)** chemokine C-X-C motif ligand-1 (CXCL-1) are not elevated in the serum of *MK2/3*-DKO mice compared to WT mice. n=6 mice/group, whereby one sample/group was pooled from 3 mouse sera. **(I)** Western Blot of immortalized WT BMDMs (iWT), immortalized *MK2*-KO BMDMs (i*MK2*-KO), and i*MK2*-KO transduced with either an empty vector as control (i*MK2*-KO+*GFP*) or rescued with *MK2* (i*MK2*-KO+*MK2*). **(J)** *Tnf* mRNA is neither affected in resting BMDM (left) nor i*MK2*-KO cells (right). Mean ± SEM, students t-test, * P<0.05, ** P<0.01, *** P<0.001.

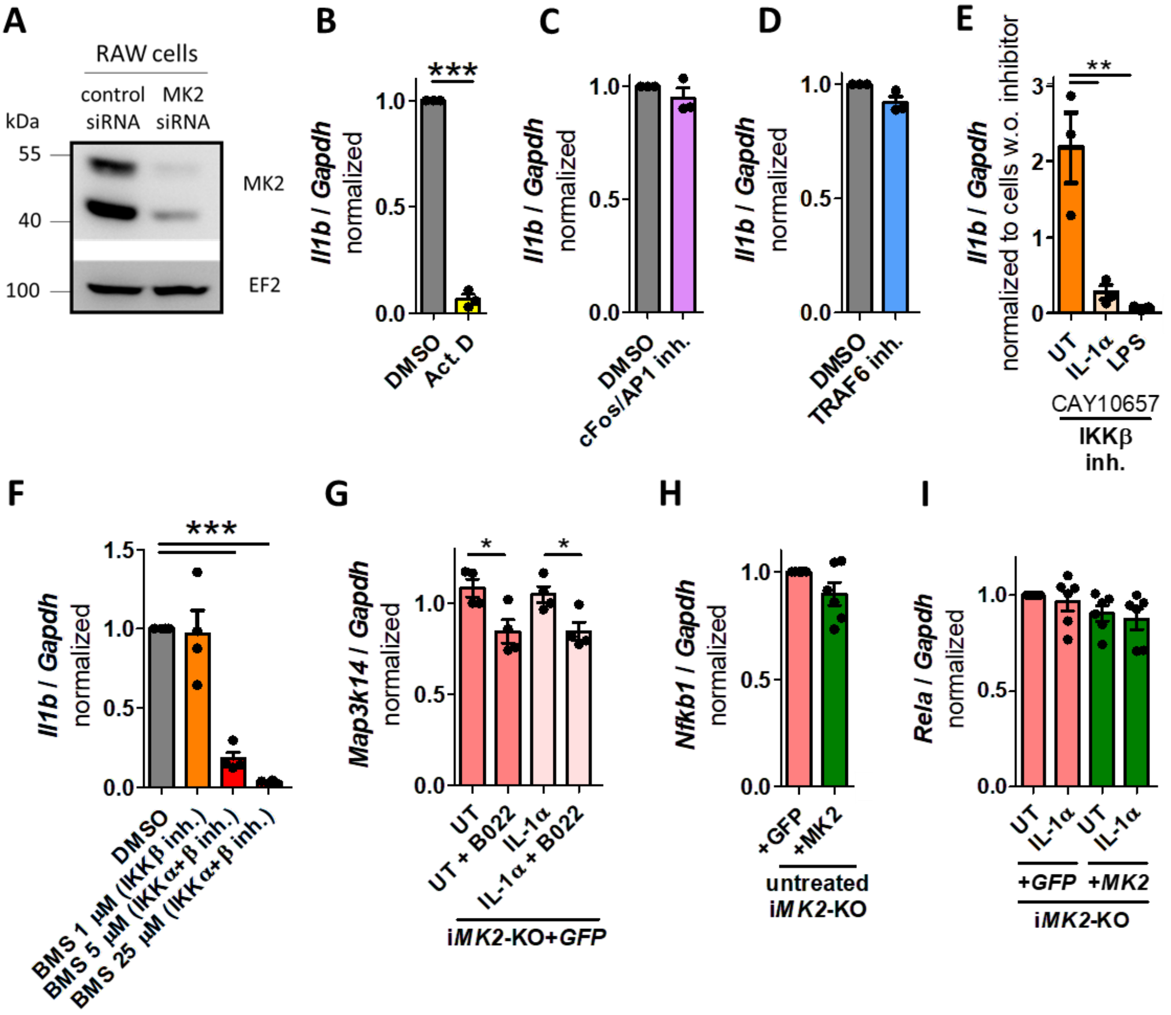


**Supplement Figure 2:**

**(A)** Western Blot of RAW cells treated either with control or *MK2* siRNA. *Il1b* mRNA levels in i*MK2*-KO+*GFP* cells after treatment with **(B)** Actinomycin D (Act. D) (10 µg/ml, 2h), **(C)** the cFos/AP1 inhibitor T-5224 (20 µM, 2h), **(D)** the SML1160 CD40-TRAF6 inhibitor (1 µM, 2h), **(E)** the IKKβ inhibitor CAY10657 (10 µM, 2h); untreated (UT), IL-1α (5ng/ml, 1h), or LPS (100 ng/ml, 1h) treated, and **(F)** IKK inhibitor BMS 345541 (2h). **(G)** *Map3k14* mRNA level after treatment with B022 inhibitor (5 µM, 2h), **(H)** *Nfkb1*, and **(I)** *Rela* mRNA levels in UT- or IL-1α (5ng/ml, 1h)-treated i*MK2*-KO+*GFP* cells compared to i*MK2*-KO+*MK2* cells. (B-D, G-I) students t-test, **(E)** 1W-ANOVA with Tukey's Multiple Comparison Test, **(F)** 1W-ANOVA with Dunnet's Multiple comparison Test, mean ± SEM, * P<0.05, *** P<0.001.

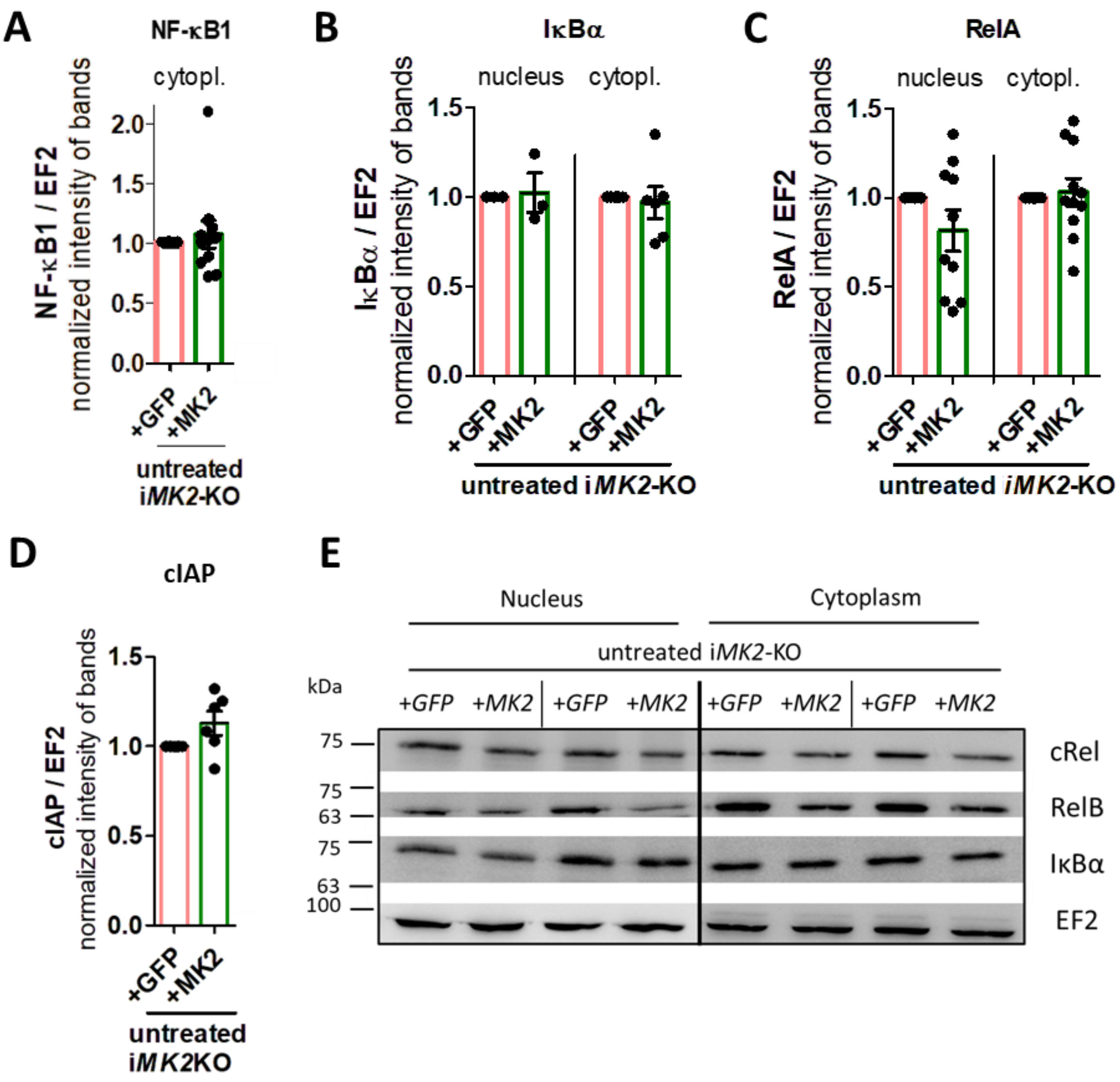


**Supplement Figure 3:**

Normalized western blot band intensities of **(A)** NF-κB1, **(B)** IκBα and **(C)** RelA in the nucleus and cytoplasm of untreated i*MK2*-KO+*GFP* and i*MK2*-KO+*MK2* macrophages. **(D)** Normalized western blot band intensities of basal cIAP of i*MK2*-KO+*GFP* and +*MK2* cells. **(E)** One representative Western Blot showing two independent experiments regarding the protein levels of cRel, RelB, IκBα and EF2 (used as a control) in the nuclear and cytoplasmic fractions of untreated i*MK2*-KO+*GFP* and +*MK2* cells. students t-test, Mean ± SEM.

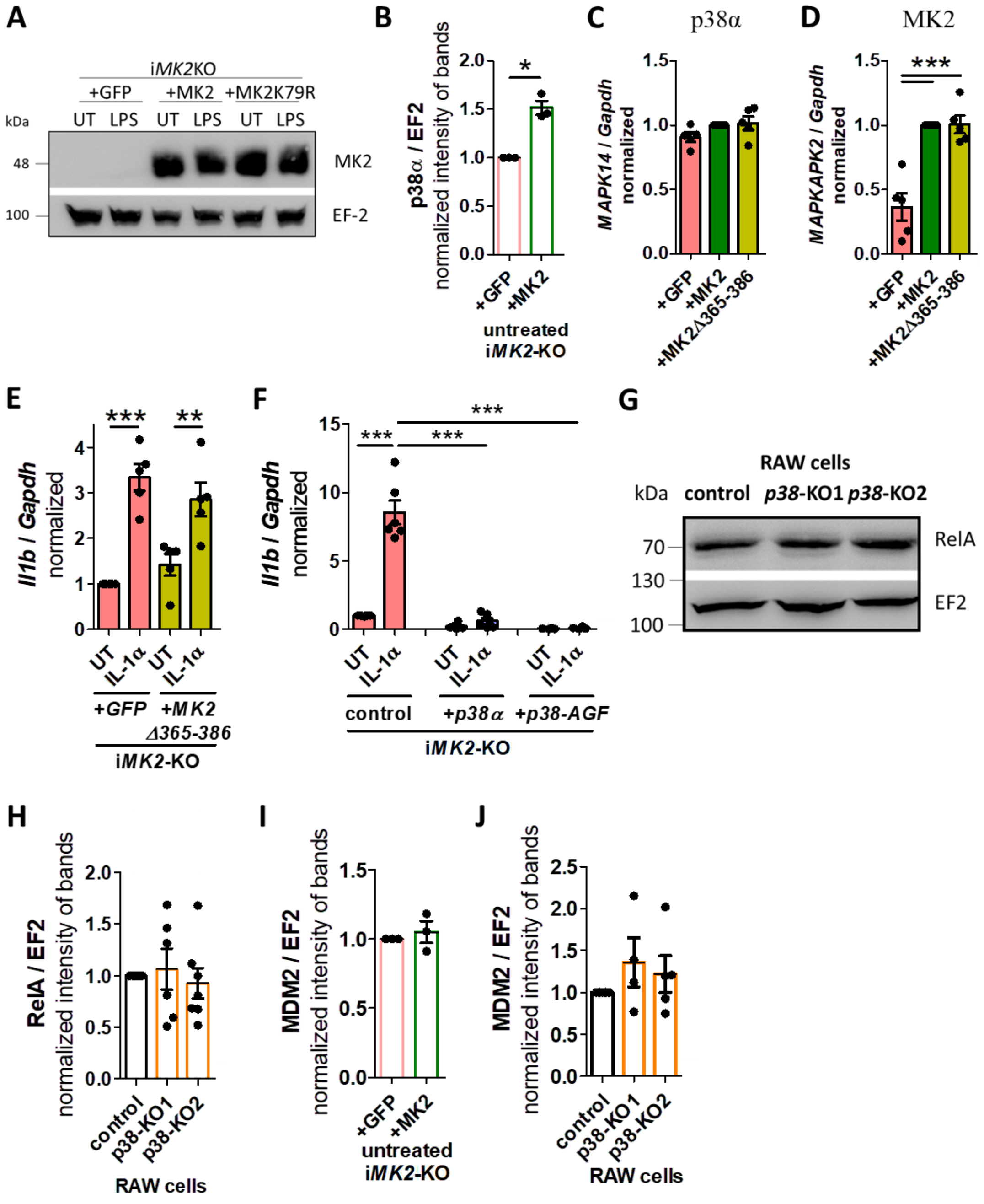


**Supplement Figure 4:**

**(A)** Rescued *MK2* and *MK2K79R* in i*MK2*-KO cells have comparable protein level. **(B)** Normalized Western Blot band intensities of p38α in untreated i*MK2*-KO+*GFP* and +*MK2* cells. **(C)** *MAPK14* and **(D)** *MAPKAPK2* mRNA levels in i*MK2*-KO+*GFP*, +*MK2*, and +*MK2-Δ365-386* cells. **(E)** MK2-Δ365-386 does not affect the *Il1b* mRNA levels in UT or IL-1α-treated cells. **(F)** *Il1b* mRNA is reduced in IL-1α (5 ng/ml, 1h)-treated i*MK2*-KO+*p38α* and kinase-inactive mutant +*p38-AGF* macrophages compared to control cells. **(G-H)** The RelA protein level is similar in *p38α*-KO cells compared to control RAW 264.1 cells. **(I)** MDM2 protein levels are similar in i*MK2*-KO+*GFP* and +*MK2* macrophages, as well as in **(J)** *p38α*-KO RAW 264.1 cells. **B, I)** students t-test, **C-J)** 1W-ANOVA followed by Tukey‘s Multiple Comparison Test, Mean ± SEM, * P<0.05, ** P<0.01, *** P<0.001.

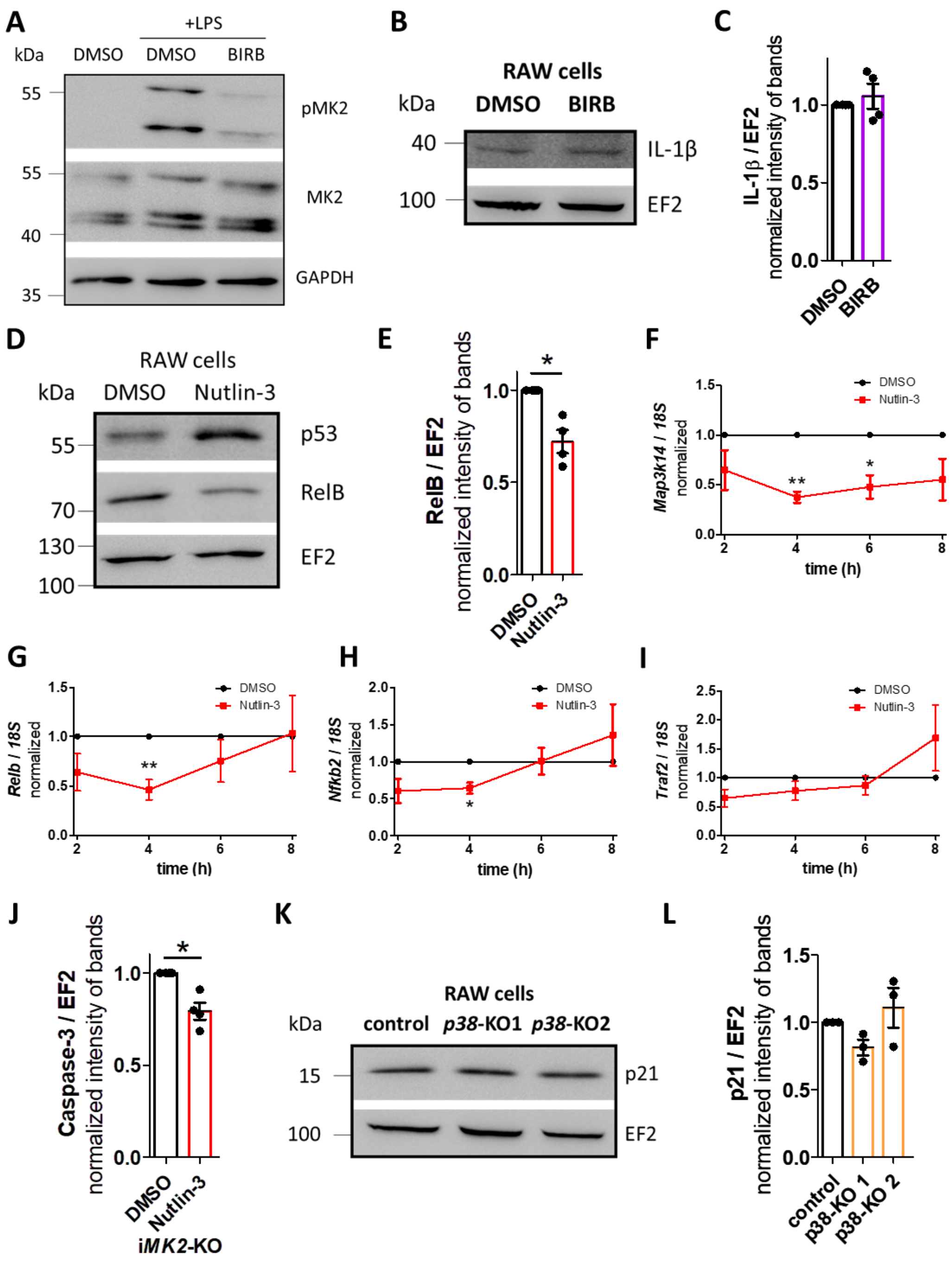


**Supplement Figure 5:**

**(A)** p38 inhibitor BIRB 796 (1 µM, 5h) inhibits phospho-MK2 (pMK2) in LPS-treated (100 nM, 30min) RAW 264.1 cells. One representative Western Blot of n=2. **(B-C)** BIRB796 (1 µM, 5h) has no effect on the level of IL-1β protein in the cytoplasmic fraction of resting RAW 264.1 cells. **(D-E)** Nutlin-3 treated (20 µM, 8h) RAW 264.1 macrophages show increased p53 and reduced RelB protein levels. **(F)** *Map3k14* (n=4), **(G)** *Relb* (n=5), **(H)** *NFkb2* (n=4) and **(I)** *Traf2* (n=5) mRNA level of Nutlin-3 (20 µM) treated i*MK2*-KO cells. **(J)** Nutlin-3 (20 µM, 4h) treated i*MK2*-KO cells have reduced levels of full-length Pro-caspase-3 protein. **(K-L)** p21 protein level in the nuclear fraction of resting *p38*-KO and control RAW 264.1 cells. **E, J)** students t-test, **F-I)** 2W-RM-ANOVA with Bonferroni posttests, mean ± SEM, * P<0.05, ** P<0.01.
